# An extended wave of global mRNA deadenylation sets up a switch in translation regulation across the mammalian oocyte-to-embryo transition

**DOI:** 10.1101/2023.03.21.533564

**Authors:** Katherine Lee, Kyucheol Cho, Robert Morey, Heidi Cook-Andersen

**Affiliations:** Department of Obstetrics, Gynecology and Reproductive Sciences, School of Medicine, University of California, San Diego, La Jolla, California, 92093, USA; Department of Molecular Biology, University of California, San Diego, La Jolla, California, 92093, USA; Department of Pathology, School of Medicine, University of California, San Diego, La Jolla, California, 92093, USA

## Abstract

The oocyte-to-embryo transition (OET) occurs in the absence of new transcription and relies on post-transcriptional gene regulation, including translational control by mRNA poly(A) tail regulation, where cytoplasmic polyadenylation activates translation and deadenylation leads to translational repression and decay. However, how the transcriptome-wide landscape of mRNA poly(A) tails shapes translation across the OET in mammals remains unknown. Here, we performed long-read RNA sequencing to uncover poly(A) tail lengths and mRNA abundance transcriptome-wide in mice across five stages of development from oocyte to embryo. Integrating these data with recently published ribosome profiling data, we demonstrate that poly(A) tail length is coupled to translational efficiency across the entire OET. We uncover an extended wave of global deadenylation during fertilization, which sets up a switch in translation control between the oocyte and embryo. In the oocyte, short-tailed maternal mRNAs that resist deadenylation in the oocyte are translationally activated, whereas large groups of mRNAs deadenylated without decay in the oocyte are later readenylated to drive translation activation in the early embryo. Our findings provide an important resource and insight into the mechanisms by which cytoplasmic polyadenylation and deadenylation dynamically shape poly(A) tail length in a stage-specific manner to orchestrate development from oocyte to embryo in mammals.

## Introduction

The transition from a fully differentiated oocyte to a totipotent embryo capable of driving development of an entirely new organism is one of the most dynamic transitions in biology. Adding to the complexity of development during these early stages, the oocyte-to-embryo transition (OET) occurs in the absence of *de novo* transcription. While the timing of developmental events differs among organisms, this period of transcriptional silence during the OET is a highly conserved phenomenon, observed from worms to humans^1–3^. However, despite the critical importance of these earliest stages of life, the molecular mechanisms required to precisely orchestrate gene expression and drive development and reprogramming across this transition in the absence of transcription remain poorly understood.

In mammals, the OET begins with global transcriptional silencing in the fully grown germinal vesicle (GV) oocyte and ends with full reactivation of transcription in the early embryo at the time of major embryonic genome activation (major EGA). Many critical developmental events occur during this window, including oocyte maturation (meiosis I), fertilization (meiosis II), reprogramming to totipotency, the first mitotic embryo cleavage, and finally, reactivation of transcription in the newly formed embryo^2,3^ (Fig. 1A). This progression of developmental events is similar in mice and humans with the exception that full transcriptional reactivation (major EGA) occurs in the 2-cell embryo in mice and even later in humans, between the 4- to 8-cell embryo stages, although increasingly sensitive approaches in recent years have uncovered low levels of transcription at the zygote (1-cell) embryo stage (minor EGA) for both mice and humans^4–6^. Proper reactivation of transcription at EGA, which marks the shift from primarily maternal to embryonic control of development, is required for development beyond the 2-cell stage in mice and the 8-cell stage in humans. The post-transcriptional mechanisms that drive gene expression and development across the OET are poorly understood, particularly in mammals where the limited number of cells available for analysis has restricted advances. However, a better understanding of these mechanisms is critical both to improve both our understanding of the earliest stages of mammalian development and the diagnosis and treatment of infertility, as only ∼50% of *in vitro* fertilized human embryos are able to successfully complete the OET, with embryo arrest commonly observed at the time EGA is to occur^7^.

**Figure 1.**
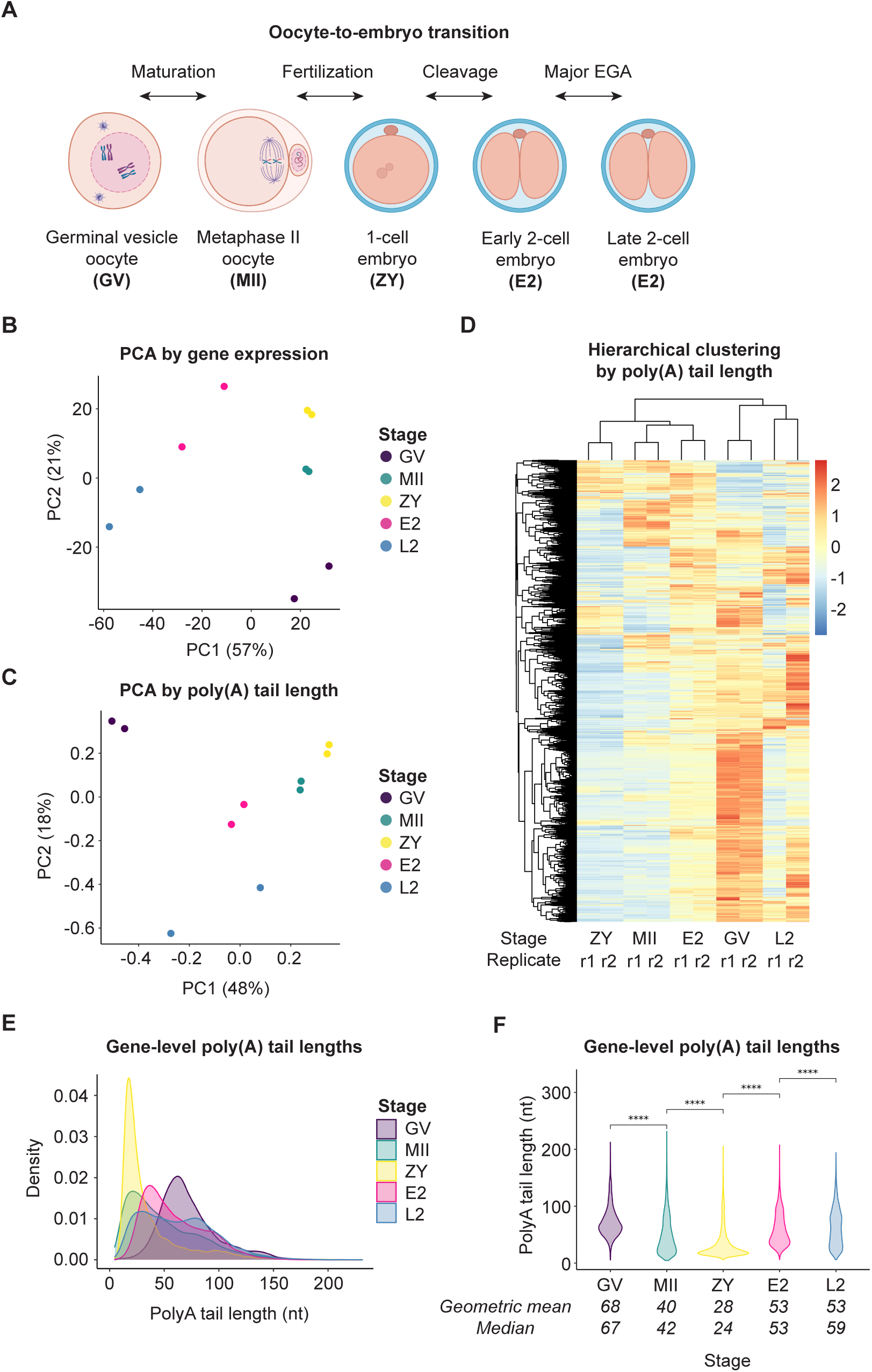
mRNA poly(A) tails are dynamically regulated across the oocyte-to-embryo transition. **(A)** Schematic of developmental stages and transitions profiled by Nanopore long-read PCR-cDNA sequencing. **(B)** PCA clustering of developmental stages and replicates by gene expression. **(C)** PCA clustering of developmental stages and replicates by poly(A) tail length. **(D)** Hierarchical clustering of developmental stages and replicates by poly(A) tail length. **(E)** Density plots showing global distributions of gene-level mean poly(A) tail lengths at each developmental stage. **(F)** Violin plots showing global distributions of gene-level mean poly(A) tail lengths at each developmental stage. Pairwise two-sided Wilcoxon tests shown each consecutive stage transition (****, p <= 0.0001). Geometric mean and median of distribution are provided below each violin plot. Only genes with a minimum of 10 polyadenylated reads in both replicates combined were included in these analyses.

Without new transcription, regulation of gene expression to drive development during the OET depends on post-transcriptional mechanisms. One mechanism known for decades to play a critical role is regulation of mRNA polyadenosine (poly(A)) tail length in the cytoplasm to regulate translation^8^. The poly(A) tail activates mRNA translation through binding by poly(A) binding protein (PABP), which, together with cap binding proteins, acts to recruit ribosomes and facilitate translation initiation^9^. However, it was recognized almost 60 years ago that sea urchin oocytes accumulate mRNAs that are not translated during oocyte growth but, instead, are stored and only translationally activated later during oocyte maturation or after fertilization^10,11^. In a series of landmark discoveries in Xenopus more than twenty years later, it was found that “dormant” oocyte mRNAs are deadenylated in the cytoplasm, stored in an unusually stable state with short poly(A) tails in the growing oocyte, and activated for translation later during the OET by cytoplasmic polyadenylation to lengthen the tail again^10–14^. These and other studies over the last 30 years have demonstrated a highly conserved role for cytoplasmic polyadenylation in the stage-specific activation of gene expression during the OET. Although these mechanisms are relatively specific to the oocyte and early embryo, similar post-transcriptional gene regulation by changes poly(A) tail length have been shown more recently to play important roles in somatic cells as well, including in cell cycle and neuronal plasticity and memory^15–18^.

In addition to cytoplasmic polyadenylation, tail shortening by cytoplasmic deadenylation is also a critical regulator of gene expression across the OET, with demonstrated roles in the repression of gene expression—both with and without decay. In somatic cells, deadenylation is the first step in many RNA decay pathways, with short tails triggering rapid decay through mRNA decapping and degradation of the body of the transcript^19,20^. In mammals, waves of deadenylation and decay occur both during oocyte maturation and before major EGA such that the majority of maternal mRNA is cleared before full reactivation of the embryonic genome^2,21–23^. In contrast, deadenylation without decay is important in the growing oocyte to translationally repress the subset of “dormant” maternal mRNAs^14^. In addition, it was posited more than two decades ago that deadenylation and decay can also be uncoupled during oocyte maturation^24–26^. Early estimations suggested that only half of the 50% decrease in polyadenylated RNA during mouse oocyte maturation resulted from RNA decay whereas the other half was likely due to deadenylation without decay. A decade later, uncoupling of deadenylation and decay was demonstrated more rigorously during maturation of Xenopus oocytes and following fertilization in the early Xenopus embryo^25,26^. In fact, mRNAs deadenylated after fertilization remained remarkably stable until the mid-blastula transition, at which point all deadenylated mRNAs were rapidly and synchronously degraded^26^. However, which specific mRNAs are regulated in this manner and the biological role for this uncoupling in development during the OET remains unknown.

While early studies of cytoplasmic polyadenylation and deadenylation across the OET were limited to single gene studies^24,27–34^, the next significant advances require the ability to examine poly(A) tail lengths and ribosome dynamics genome-wide to uncover patterns and identify regulatory principles. Within the past decade, approaches to capture and accurately determine the length of the homopolymeric poly(A) tail genome-wide have been developed^16,17,35^, and increasingly sensitive approaches that allow accurate tail length determination in the small number of oocytes and embryos that can be feasibly obtained in mammalian systems have rapidly emerged in recent years^36–41^. These approaches have enabled genome-wide pictures of maternal mRNA tail dynamics across the OET in oocytes and embryos in frogs, zebrafish, flies, and, very recently, in humans^16,35,42–44^, with studies in progress for mice, rats, and pigs^45,46^. The newest protocols are long-read sequencing approaches, which, in contrast to early short-read approaches, can capture and sequence even very long poly(A) tails at isoform-level resolution. Importantly, parallel advances in the sensitivity of ribosome profiling have also allowed demonstration of the coupling between poly(A) tail regulation and ribosome association in the oocyte and early embryo in an evolutionarily conserved manner^16,32,43,47^. However, the transcriptome-wide landscape of mRNA poly(A) tail lengths across the entire OET in mice—and how changes in poly(A) tails affect translational efficiency across the OET in mammals—remains unknown.

Here, we leverage a new Nanopore cDNA-PCR long-read sequencing approach to determine and compare poly(A) tail lengths and mRNA abundance genome-wide in mice across five stages of development from oocyte to embryo. Integration of these data with recently published ribosome profiling data at the same developmental stages^47^ allowed us to examine patterns of regulation between mRNA tail poly(A) tail length and translational efficiency across the entire OET for the first time in a mammalian system. We identify global and gene-specific patterns in tail length dynamics driving gene expression regulation during oocyte maturation, fertilization, and embryonic genome activation. Together, these data provide a rich resource for exploring the mechanisms that control gene expression at multiple levels and provide fresh insights into the relationship between poly(A) tail regulation and translation across the OET in mammals.

## Results

### Determination of poly(A) tail lengths genome-wide across the mammalian oocyte-to-embryo transition

To determine and compare mRNA tail lengths transcriptome-wide at each stage across this transition in a small number of cells, we tested the accuracy, sensitivity, and reproducibility of tail lengths determined using a newly developed Nanopore cDNA-PCR protocol, together with a published algorithm able to determine the length of homopolymeric polyadenosine sequences in Nanopore raw data^48^. This long-read sequencing protocol involves ligation of an adapter to mRNA 3’ ends before reverse transcription, which allows capture and preservation of mRNAs with poly(A) tails as short as 10 nucleotides and the opportunity to simultaneously determine mRNA tail length and abundance (Fig. S1A). To examine the accuracy of tail length determinations, we sequenced cDNA standards of known poly(A) tails ranging from 10 to 150 nucleotides (nt) and found that this approach was able to accurately determine mRNA tail lengths within this range and to distinguish between tails with only 10 nucleotides difference in length (e.g., 40 vs. 50 nt) (Fig. S1B). Tail length determinations did not differ by read type (Fig. S1C) and were highly reproducible between replicates (Fig. S1D). To test the sensitivity of the protocol with low RNA input, we examined mRNA tail lengths and RNA abundance in Hela cells with total RNA input as low as 200 ng. With this input, we obtained high genome coverage, with 13,727 genes represented by polyadenylated reads. Further, the mean gene-level mRNA tail length was 94 nucleotides (Fig. S1E), which is within the range of poly(A) tail lengths previously reported in Hela^16,17,41,49^. Together, these data demonstrate that this long read Nanopore approach can determine poly(A) tail lengths accurately and reproducibly genome-wide with relatively low RNA input.

We next applied this approach to sequence and simultaneously determine poly(A) tail lengths and mRNA abundance genome-wide at five developmental stages across the oocyte-to-embryo transition: the fully grown germinal vesicle oocyte (GV), MII oocyte (MII), zygote (ZY), early 2-cell embryo (E2) and late 2-cell embryo (L2) (Fig. 1A). These five stages span development from the fully grown, transcriptionally silent oocyte to after transcription reactivation at major EGA in the 2-cell embryo. At each stage, two biological replicates were sequenced with ∼400 oocytes or embryos each. We obtained high genome coverage, with an average of 12,204 genes represented by all polyadenylated reads in both replicates at each developmental stage and 8,490 genes represented by at least 10 polyadenylated reads in both replicates (Data S1). Principle component analysis (PCA) of normalized gene expression showed that the biological replicates for each stage clustered together (Fig. 1B). In contrast, transcriptome profiles for the different developmental stages were distinct and well separated overall, with the stages before (GV, MII, ZY) and after (E2, L2) embryonic genome activation being more closely clustered with each other, as expected. Following normalization, read counts for a panel of control RNAs covering a wide range of known lengths and concentrations spiked into each RNAseq reaction^50,51^ correlated well with input concentrations within each stage and replicate (Fig. S2A). With respect to poly(A) tail length, the biological replicates for each stage also clustered together both by PCA (Fig. 1C) and hierarchical clustering (Fig. 1D) although tail length patterns are more variable at L2, likely because many maternal mRNAs are being degraded and replaced with new embryonic transcripts at this stage. In contrast to trends for the transcriptome, the tail lengths for the two most temporally separated stages—GV and L2—were more similar to each other than to the intervening stages, which clustered together. Overall, replicates for each stage were highly reproducible, with high correlations in both measured mRNA abundance and poly(A) tail length (Figs. S2B-C).

Comparison of results across consecutive stages allowed us to determine changes in tail length and mRNA abundance across four central developmental transitions—oocyte maturation (GV to MII), fertilization (MII to ZY), the first mitotic cleavage and minor EGA (ZY to E2), and major EGA (E2 to L2) (Fig. 1A). To validate detected changes in poly(A) tail length, we examined changes in mRNA poly(A) tail length for a subset of genes previously shown to be regulated by poly(A) tail length during oocyte maturation (Figs. S3A-B)^24,27–34^. For example, *Btg4, Ccnb1, Mos,* and *Tex19.1* have been previously shown to be polyadenylated during oocyte maturation, and we also find that each of these mRNAs demonstrate a significant increase in poly(A) tail length from GV to MII. Similarly, *Actin, α-tubulin*, *Smc4*, and *Zp2* are known to be deadenylated during oocyte maturation, and we observe a significant decrease in tail length for each of these mRNAs from GV to MII^24,33^. Together, these data provide a robust validation of our approach to determine and compare poly(A) tail lengths genome-wide within and between each developmental transition from oocyte to embryo.

### Poly(A) tail lengths are globally regulated in a stage-specific manner

We first examined global trends in tail length regulation, comparing the global distribution of poly(A) tail lengths at each stage at the gene level. Global mean poly(A) tail lengths were longest at the GV stage and significantly longer at this stage relative to all other stages examined, with a mean of 68 nucleotides (Figs. 1E-F). Poly(A) tail lengths then globally shorten during oocyte maturation to a mean of 40 nucleotides in MII oocytes, consistent with a wave of mRNA deadenylation and decay previously described during oocyte maturation^23,24^. Unexpectedly, however, we found that global poly(A) tail lengths are significantly shortened even further in an extended wave of global deadenylation during fertilization. In fact, mean poly(A) tail lengths at the zygote stage are significantly shorter than at any other stage across the oocyte-to-embryo transition, with a mean length of only 28 nucleotides and only ∼5% of genes with their longest mean tail lengths at the zygote stage. This overall global trend of progressive poly(A) tail shortening in the oocyte and early embryo is consistent with the eventual decay of the majority of the maternal mRNA pool prior to major EGA in the 2-cell embryo as part of the maternal-to-zygotic transition^21,22^.

Following the increasingly short global tail lengths observed during oocyte maturation and fertilization, tails globally lengthen again during the first mitotic cleavage cycle from ZY to E2 and from E2 to L2, with a mean length of 53 nucleotides at both the E2 and L2 stages (Figs. 1E-F). Lengthening of poly(A) tails during these stages might represent polyadenylation of existing maternal transcripts, new transcription of the same genes in the embryo, or a combination of these mechanisms. Temporally, tail lengthening at E2 and L2 coincides with reactivation of transcription in the new embryo, and this relationship appears to be evolutionarily conserved as others have observed a similar trend at EGA in zebrafish and Xenopus embryos^16,42^. That tail lengthening might be driven, at least in part, by new transcription is also consistent with published reports in somatic cells that show that newly transcribed mRNAs typically have longer tails on average relative to older cytoplasmic mRNAs^52–54^. Together, these data demonstrate that mRNA tail lengths across the oocyte-to-embryo transition are not random but are, instead, highly regulated at the global level in a stage-specific manner. A resource containing measured poly(A) tail lengths for all genes at each developmental stage is provided (Data S2).

### Poly(A) tail lengths are selectively regulated at the gene-specific level

The current model for post-transcriptional regulation of the maternal mRNA pool would predict that mRNAs with important roles in development will be selectively polyadenylated and/or deadenylated in a stage-specific manner to regulate translational activation and repression, respectively, irrespective of the global trend at that stage transition. Therefore, we next asked whether, amid these dynamic global changes, the poly(A) tails of specific subsets of mRNAs are selectively regulated at a gene-specific level. To address this question, we compared poly(A) tail lengths for each gene across each of the four developmental transitions (see Methods). Consistent with the global trend of tail shortening during oocyte maturation, we found that mRNAs for approximately two-thirds of genes are significantly deadenylated from GV to MII. However, the remaining one-third of genes escape this global wave of deadenylation, having tail lengths that are either not significantly changed or significantly polyadenylated (Figs. 2A-B). Similarly, during fertilization, although the global mean poly(A) tail length is significantly further decreased (Figs. 1E-F), more than half of the genes escape this global trend, with tail lengths that are either not significantly changed or significantly increased between MII and ZY (Figs. 2A-B).

**Figure 2.**
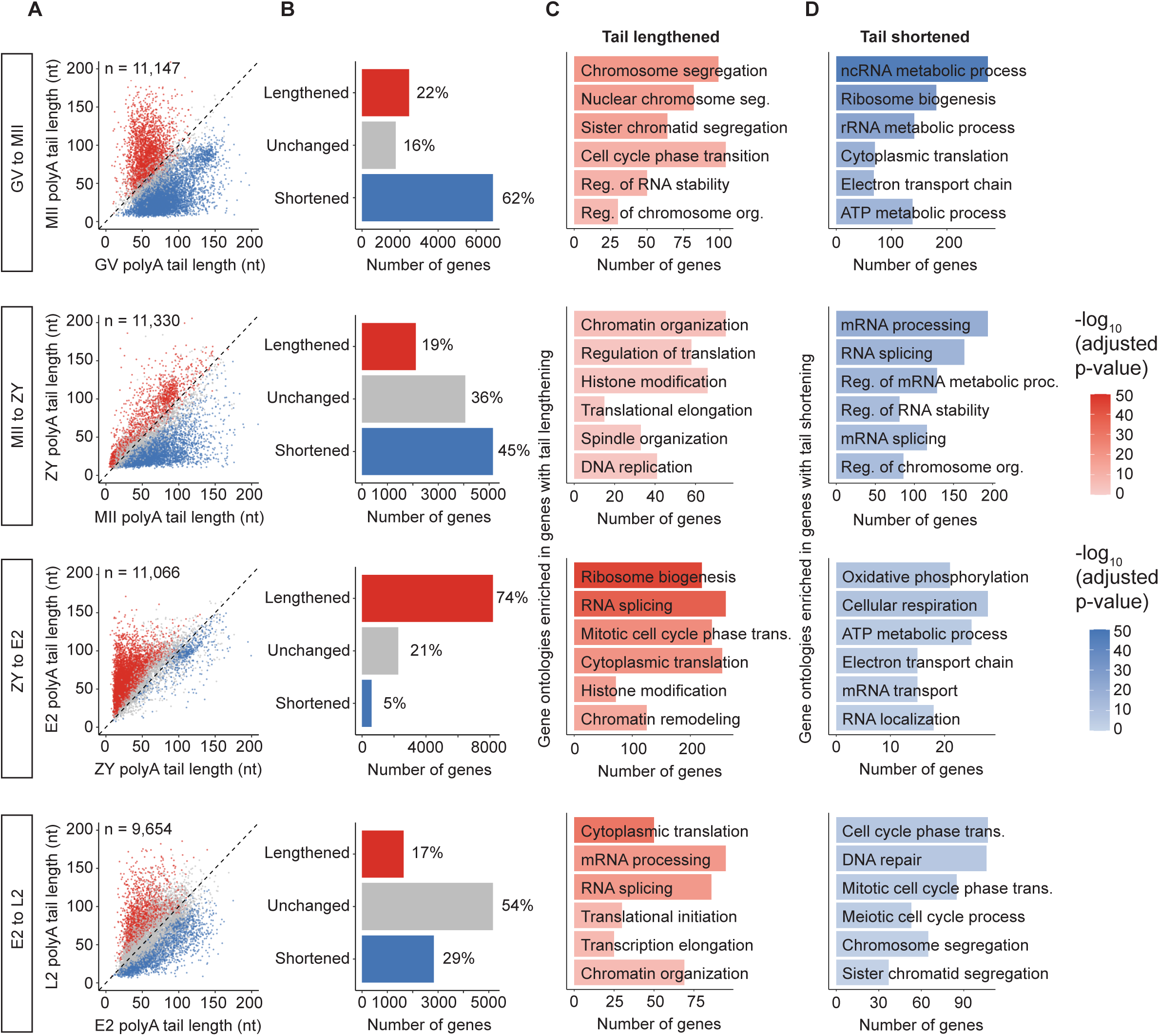
mRNA poly(A) tail lengths are regulated in a gene-specific manner. **(A)** Scatter plots showing geometric mean poly(A) tail lengths for genes with significantly increased (lengthened, red), decreased (shortened, blue) or unchanged (grey) tail length at each consecutive developmental stage transition (adjusted p-value < 0.05). Only genes with at least 10 polyadenylated reads in both stages represented were included in this analysis. **(B)** Number of genes in each category shown in **(A)**. **(C-D)** Gene ontologies enriched in genes with lengthened (red) **(C)** or shortened (blue) **(D)** tail lengths at each stage transition.

Gene-specific regulation in the opposite direction is also seen at stages for which the global trend is an increase in mean tail length. During the transition from ZY to E2, the global trend undergoes a striking shift from tail shortening to tail lengthening, with approximately three-quarters of genes being significantly polyadenylated. The remaining one-quarter of genes have poly(A) tail lengths that are either not significantly changed or significantly decreased (Figs. 2A-B). From E2 to L2, while the increase in tail length is modest at the global level (Figs. 1E-F), examination at the gene level reveals dynamic changes in tail length across this transition, with almost half of all genes undergoing significant increases or decreases in tail length from E2 to L2 (Figs. 2A-B). These opposing trends are consistent with both the widespread clearance of maternal mRNA and resumption of transcription during major EGA. Together, these data demonstrate that, despite the striking global changes in poly(A) tail length observed, a substantial proportion of mRNAs are protected from—or even regulated in a manner opposite to–the global trend at each developmental transition. These findings raise the possibility that this gene-specific regulation of poly(A) tails plays an important role in the selective activation and repression of genes with important roles in regulating developmental progression at each stage. A resource containing changes in poly(A) length for each gene across each stage transition is provided (Data S3).

### mRNAs with gene-specific changes in poly(A) tail length are enriched for stage-specific roles in development

If gene-specific changes in poly(A) tail length are important to regulate expression of genes needed to drive development from oocyte to embryo, one would predict that mRNAs with significant changes in tail length are enriched for genes with important developmental roles at each stage. To determine whether this is the case, we conducted gene ontology (GO) overrepresentation analysis to identify gene sets enriched in mRNAs with significant changes in poly(A) tail length across each stage transition (Figs. 2C-D; Data S4). During oocyte maturation from GV to MII, mRNAs with significant increases in tail length were enriched for genes with biological processes related to meiosis (“chromosome segregation”, “sister chromatid segregation”, and “cell cycle phase transition”), consistent with progression from prophase I to metaphase II across this transition. During fertilization from MII to ZY, mRNAs with tails that significantly increased were highly enriched for roles related to epigenetic modification (“chromatin organization”, “histone modification”), replication (“DNA replication”), and translation (“regulation of translation”, “translational elongation”), consistent with reprogramming to totipotency, preparation for the first mitotic cleavage cycle, and a known increase in protein synthesis from the MII to ZY stages^55^.

In the early embryo from ZY to E2, genes with significant tail lengthening were enriched for processes related to mRNA production (“RNA splicing”), translation (“ribosome biogenesis”, “cytoplasmic translation”), mitosis (“mitotic cell cycle phase transition”), and epigenetic modification (“chromatin remodeling”, “histone modification”). These findings are consistent with resumption of transcription in the newly formed embryo during minor EGA, the first mitotic division and continued chromatin reprogramming at this stage. From E2 to L2, mRNAs with increased tail lengths were enriched for roles in transcription (“DNA-templated transcription, initiation and elongation”, “RNA splicing”, “mRNA processing”), translation (“cytoplasmic translation,” “translational initiation”), and epigenetic modification (“chromatin organization”), consistent with the major wave of EGA, increased translational activity, and continued chromatin remodeling.

In contrast to polyadenylation, mRNAs significantly deadenylated across each stage transition were less consistently enriched for stage-specific factors in development that would be predicted to be downregulated. This is consistent with the possibility that deadenylation is a more global and less selective process (Fig. 2D). However, that genes polyadenylated at each transition were enriched for factors involved in important—and highly relevant—pathways appropriate for each stage transition suggests that polyadenylation is a specific process and identifies these genes as candidates for factors with important roles in the transition from oocyte to embryo.

### Deadenylation is uncoupled from decay in a gene-specific manner during oocyte maturation

Accumulated data suggest deadenylation is not obligatorily coupled to decay during mouse oocyte maturation^24–26^. However, previous studies were unable to determine the proportion of maternal mRNAs affected by this phenomenon or what role the deadenylated but stable mRNAs might play in development. Subsequent attempts to identify the specific mRNAs that escape decay following deadenylation during oocyte maturation in mice have been limited by technical challenges^56^, including the inability to measure poly(A) tail length and mRNA abundance simultaneously. Consequently, the prevalence of this phenomenon, the specific maternal mRNAs for which deadenylation is uncoupled from decay and the potential biological role of this process in development from oocyte to embryo remain largely unexplored.

Focusing on oocyte maturation, where this phenomenon has been previously described^24^, we identified the specific subsets of deadenylated mRNAs that are degraded versus those that escape decay by comparing changes in poly(A) tail length to changes in mRNA abundance from GV to MII. We found that, of the 6,709 genes significantly deadenylated during oocyte maturation (see Methods), 3,793 (57%) were also downregulated. Because our approach can capture mRNAs with short tails, this decrease in mRNA abundance most likely represents mRNA decay (deadenylated-decayed mRNAs). In contrast, 2,916 of the deadenylated genes (43%) demonstrated no evident downregulation but instead remained stable (deadenylated-stable mRNAs) (Fig. 3A). Consistent with previous findings^24–26^, these data further support that mRNA deadenylation and decay can be uncoupled during oocyte maturation.

**Figure 3.**
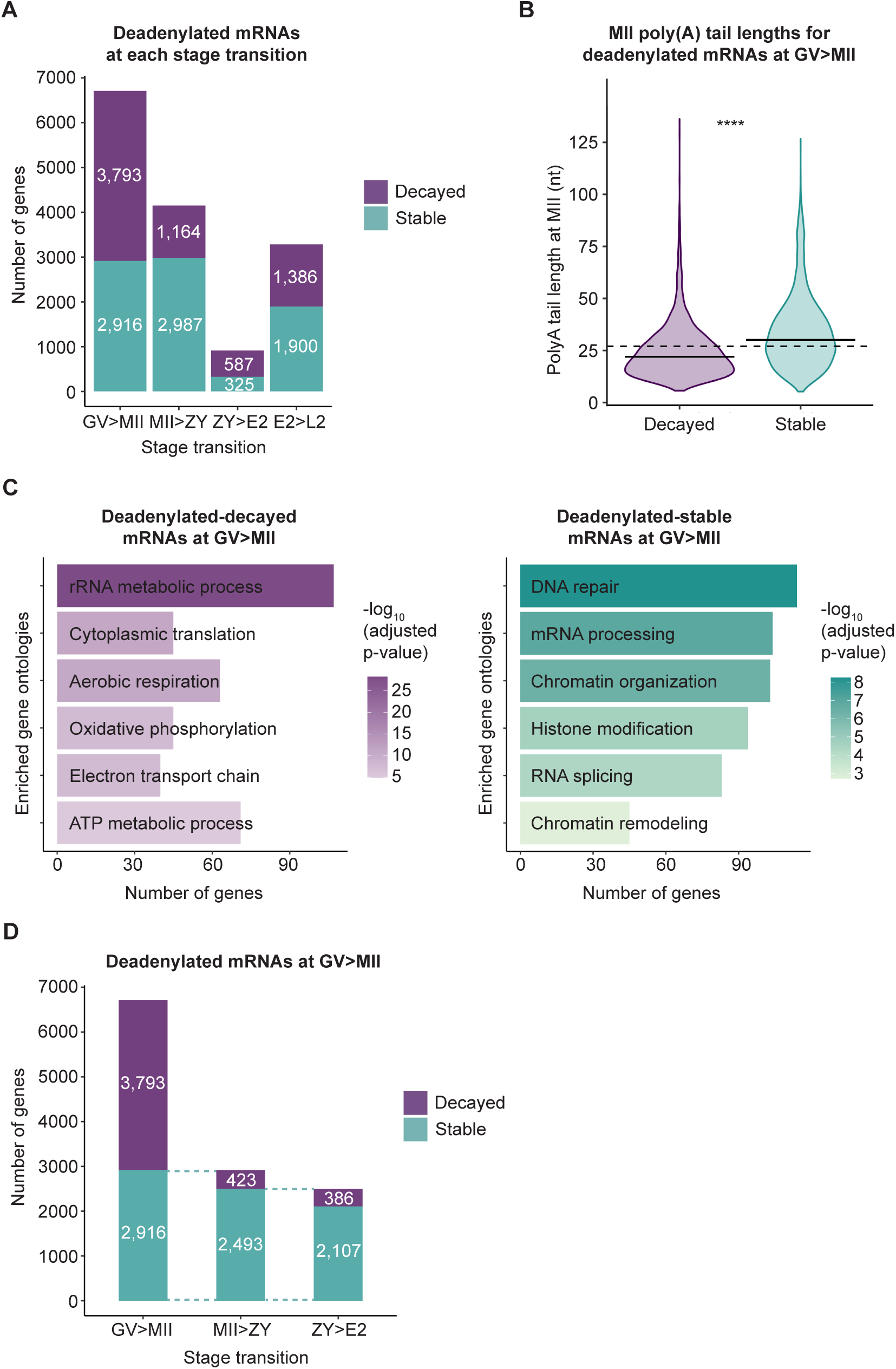
Deadenylation and decay can be uncoupled during the oocyte-to-embryo transition. **(A)** Number of deadenylated genes that are downregulated (decayed, purple) or unchanged (stable, teal) in abundance at each stage transition. **(B)** Poly(A) tail lengths at the MII stage of genes that are deadenylated and decayed or deadenylated and stable during oocyte maturation. Solid horizontal lines indicate geometric means. Dashed horizontal line at indicates predicted PABP footprint of 27 nt. Two-sided Wilcoxon test shown (****, p <= 0.0001). **(C)** Gene ontologies enriched in genes deadenylated and decayed (purple) or deadenylated and stable (teal) during oocyte maturation. **(D)** Number of genes that were deadenylated and stable from GV to MII that remain stable or are decayed from MII to ZY and from ZY to E2.

Whether uncoupling of deadenylation and decay also occurs at later stages during the oocyte-to-embryo transition remains unknown. To explore this possibility, we compared changes in poly(A) tail length to changes in mRNA abundance across each subsequent stage transition. Expression patterns of deadenylated mRNAs that were stable versus degraded were similar to those observed from GV to MII. Specifically, we identified distinct subsets of deadenylated mRNAs with and without decreases in mRNA abundance from MII to ZY, ZY to E2, and E2 to L2 (Fig. 3A), consistent with the possibility that deadenylation might also be uncoupled from decay during fertilization and in the early embryo. However, the global trend in poly(A) tail length shifts from tail shortening to lengthening at the ZY to E2 transition with fewer genes deadenylated overall (Fig. 2B), and analyses of RNA abundance at these subsequent stages, particularly from E2 to L2, are confounded by transcription at EGA although newly transcribed mRNA would be predicted to have longer, not shorter, tails^52–54^. For these reasons, we focused further analyses only on mRNAs deadenylated during oocyte maturation.

The mechanism by which the degraded and stable subsets of deadenylated mRNAs are differentially regulated is unknown and likely multi-factorial. One possibility is that the stable subset of deadenylated mRNAs, as a group, maintain significantly longer poly(A) tails than the degraded subset. Indeed, at the MII stage, we found that the mean tail lengths for mRNAs deadenylated but stable were significantly longer than mRNAs that were degraded (Fig. 3B). Of note, however, the tail length distributions for both groups were relatively short, and the magnitude of the difference in tail length was small, with a mean of 30 nucleotides in the stable subset compared to 22 nucleotides in the degraded subset. That these mean tail lengths for the stable and degraded mRNAs hover just above and below the predicted binding footprint for a single cytoplasmic poly(A) binding protein (estimated at 27 nucleotides in mammals^57^), respectively, suggests that deadenylation past the length required for a single poly(A) binding protein to bind might play a critical role in determining mRNA fate during oocyte maturation. Consistent with this idea, it has been shown that mRNAs with tails of ∼25 nucleotides or less are destabilized when PABPC levels are reduced in Hela cells^58^. Together, these data identify the specific subset of deadenylated maternal mRNAs uncoupled from decay during oocyte maturation and suggest that small differences poly(A) tail length at the MII stage might be one mechanism mediating the uncoupling of deadenylation and decay and determining mRNA fate.

### Global deadenylation primes maternal mRNAs for selective readenylation and activation in the early embryo

To explore the potential biological role of uncoupling of deadenylation and decay during global deadenylation from GV to MII, we first asked in what biological processes the degraded versus stable gene subsets are involved. GO overrepresentation analysis revealed that, among the genes enriched in the deadenylated-stable mRNAs during oocyte maturation, were factors for processes important—not in the oocyte—but for development in the zygote and later stages of embryo development (Fig. 3C). These factors included those with important roles in replication, transcription and reprogramming in the early embryo, including “DNA replication,” “DNA repair”, “chromatin organization,” “chromatin remodeling,” “histone modification,” “RNA splicing,” and “mRNA processing”. In contrast, GO categories for the deadenylated-decayed mRNAs were enriched for processes known to be downregulated during oocyte maturation^59^, such as “cytoplasmic translation,” consistent with a known decrease in translation during oocyte maturation, as well as metabolic processes (“aerobic respiration,” “oxidative phosphorylation,” “ATP metabolic process”), consistent with the maturing oocyte transitioning to a metabolically quiescent state (Fig. 3C). Enrichment of deadenylated-stable genes for roles specific to the early embryo suggests that stabilization of deadenylated mRNAs is a selective process and raises the intriguing possibility that these deadenylated-stable mRNAs may be retained for translational activation at later stages in the early embryo.

To investigate the possibility that deadenylated-stable mRNAs are activated at later stages, we next determined the fate of this subset of mRNAs following oocyte maturation at the next developmental stage—in the zygote. We found that 2,493 of 2,916 (85%) of these genes remain stable during fertilization from MII to ZY (Fig. 3D, MII>ZY), and 546 of these stable genes (22%) have tails that are significantly lengthened again during this transition (Figs. 4A-B, MII>ZY). For the majority of this subset, this tail lengthening almost certainly represents readenylation of maternal mRNAs as new transcription during this transition is relatively low, and only 118 (22%) of these mRNAs with increased tail length overlap with >4000 genes previously predicted to be minor EGA genes^4^. In the context of global tail length shortening from MII to ZY (Figs. 1E-F), this subset of readenylated mRNAs represents one quarter (26%) of all genes with significant increases in tail length from MII to ZY.

**Figure 4.**
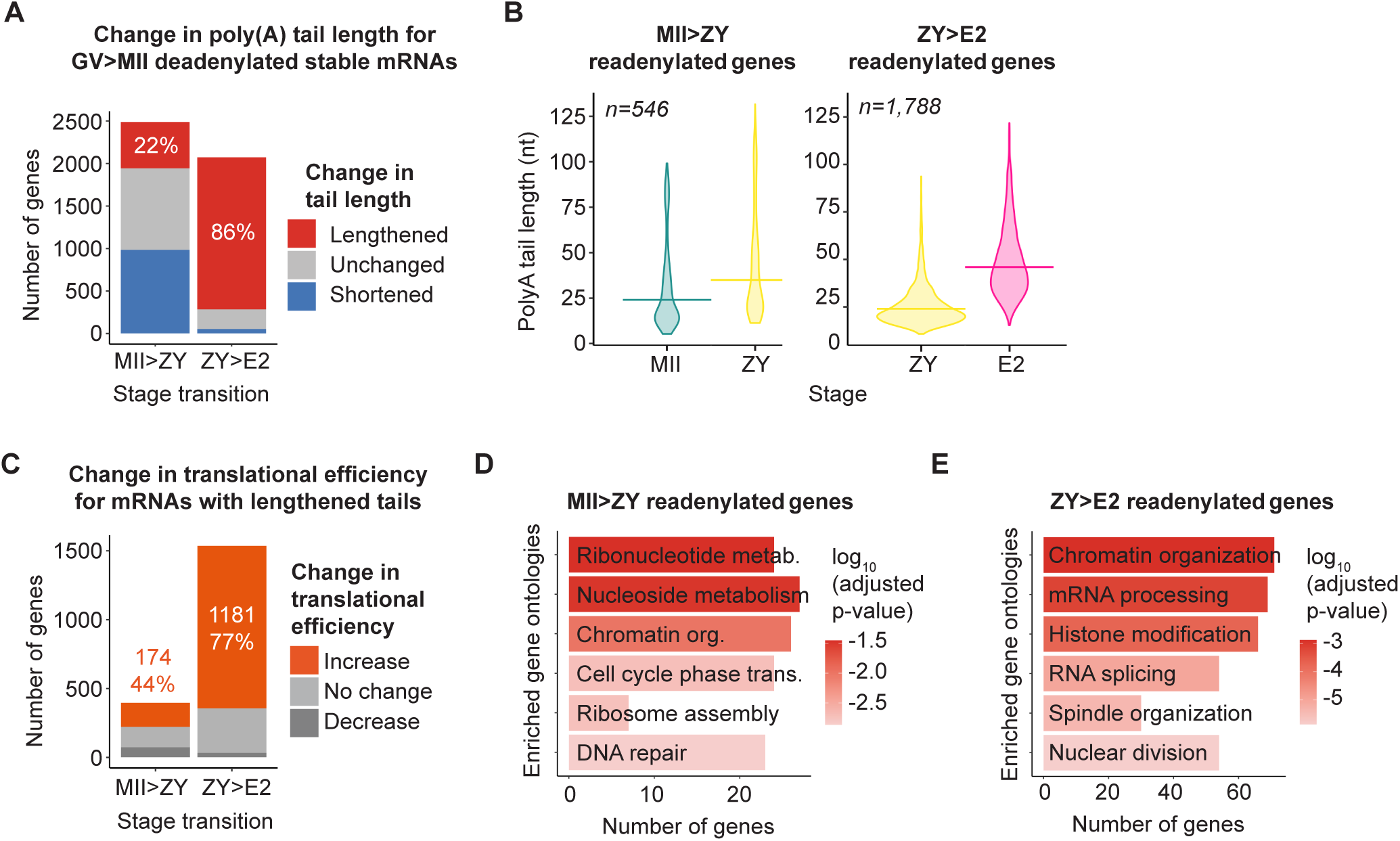
mRNA readenylation in the embryo. For **(A-C, MII>ZY)** and **(D)**, genes examined were deadenylated and stable from GV to MII and remain stable from MII to ZY. For **(A-C, ZY>E2)** and **(E)**, genes examined were deadenylated and stable from GV to MII and remain stable from MII to ZY and from ZY to E2. **(A)** Proportion of these genes with different changes in poly(A) tail length from MII to ZY (left) or ZY to E2 (right). Percentage of readenylated (lengthened) genes is indicated. **(B)** Poly(A) tail lengths at MII and ZY stages for these genes that are readenylated from MII to ZY (left) or ZY to E2 (right). Horizontal lines indicate geometric means. **(C)** Proportion of readenylated genes with different concurrent changes translational efficiency from MII to ZY (left) or ZY to E2 (right). Percentage of genes with increased translation efficiency is indicated. Genes were filtered by a minimum of 1 FPKM in the RNA-seq dataset used to calculate translational efficiency. **(D)** Enriched gene ontologies in genes readenylated during from MII to ZY. **(E)** Enriched gene ontologies in genes readenylated during from ZY to E2.

If readenylation of this subset of mRNAs is biologically important, one would predict that the readenylated mRNAs are translated and that they play relevant roles in development. With respect to translation, analysis of a previously published ribosome profiling dataset^47^ revealed that the vast majority of these readenylated mRNAs (75%) are associated with ribosomes in the zygote and almost half (44%) of those for which translational efficiency could be calculated demonstrate an increase in translational efficiency between MII and ZY stages (Fig. 4C, MII>ZY). Consistent with important roles for these mRNAs in the newly fertilized embryo, this readenylated subset of genes remains enriched for factors important for early embryo development. These processes include “ribonucleotide metabolic process” and “nucleoside phosphate metabolic process,” consistent with DNA replication and the gradual increase in transcriptional activity in the zygote; “DNA repair”, “cell cycle phase transition,” and “chromatin organization,” consistent with mitosis and reprogramming at the zygote stage; as well as “ribosome assembly,” consistent with the demonstrated increase in ribosome production and translational activity in the zygote^55,60^ (Fig. 4D).

Since a relative minority of the genes that were deadenylated but stable from GV to MII were readenylated in the zygote, but a large proportion remained stable through fertilization (2,493 genes, Fig. 3D, MII>ZY), we next investigated the fate of these genes at the next developmental transition—from ZY to E2. We found that 2,107 (85%) also remain stable during this transition (Fig. 3D, ZY>E2), and, remarkably, 1,788 (86%) of these genes have tails that are significantly lengthened during this transition, representing almost a quarter (22%) of all the genes we observe with significant tail lengthening from ZY to E2 (Figs. 4A-B, ZY>E2). That only 20% of these genes overlap with mRNAs previously predicted to be minor EGA genes^4,55^ suggests that the majority of this population represents maternal mRNAs readenylated in the 2-cell embryo and not new transcription. The vast majority of this readenylated subset are associated with ribosomes at the E2 stage (89%) and have increased translational efficiency between ZY and E2 (77%) (Fig. 4C, ZY>E2). Further, these genes are enriched for factors with important roles in “RNA splicing” and “mRNA processing”, consistent with the resumption of transcription; “chromatin organization” and “histone modification”, consistent with continued reprogramming; and “spindle organization” and “nuclear division”, consistent with the first mitotic cleavage from ZY to E2 (Fig. 4E). Together, these results support a model in thousands of maternal mRNAs can be selectively stabilized during oocyte maturation and selectively readenylated at later stages, including the transition from MII to ZY and, even more markedly, during ZY to E2, to regulate gene expression and development in the early embryo before major EGA.

### Poly(A) tail length is positively coupled with translation efficiency from oocyte to embryo

A key question is whether these changes in poly(A) tail length drive changes in translational efficiency at each stage. A positive correlation between mRNA poly(A) tail length and translational efficiency has been observed during the oocyte-to-embryo transition in Drosophila, Xenopus, and zebrafish^16,35,43^. As previously mentioned, this coupling between tail length and translational efficiency is relatively unique to the oocyte and the early embryo and is not detected in somatic cells or even at more advanced stages of development^16–18^. The timing of uncoupling varies somewhat between organisms but, overall, coupling is no longer evident by gastrulation^16,43^. In mammals, coupling between tail length and translational efficiency is also seen in the mouse GV and MII oocyte^32,47^. However, the relationship between tail length and translational efficiency during fertilization and in the early mouse embryo remains largely unexplored.

To determine the degree to which changes in poly(A) tail length represent differences in translational efficiency in the mammalian oocyte and early embryo, we integrated mRNA poly(A) tail length measured in our dataset with mRNA translational efficiency at the same stages of development across the OET as determined in a recently published Ribo-seq dataset^47^. These analyses demonstrated that, indeed, poly(A) tail length and translational efficiency are positively correlated at each of the five developmental stages from GV to L2 (Figs. 5A, S4A). Consistent with findings in other organisms^16,43^, we did not observe a clear decrease in coupling following EGA, suggesting that uncoupling of tail length and translational efficiency in mice occurs at a later stage than examined in this study. We next investigated the relationship between change in tail length and change in translational efficiency between consecutive developmental stages. As observed within each individual stage, the change in tail length and change in translational efficiency across consecutive stages were also coupled across all four developmental transitions (Figs. 5B, S4B). Together these data demonstrate that poly(A) tail length is an important determinant of translational efficiency to activate and repress gene expression in both mouse oocytes and early embryos.

**Figure 5.**
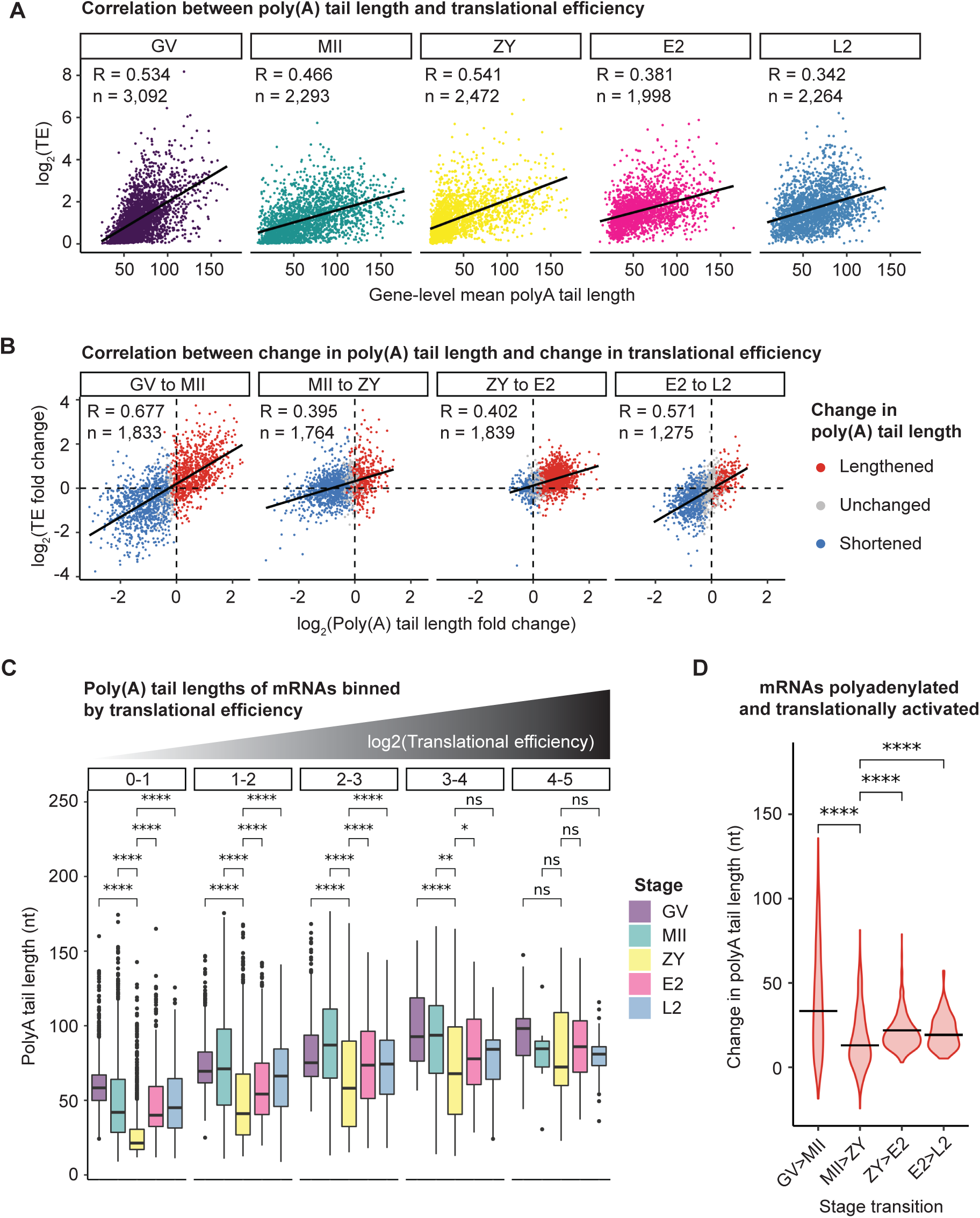
Poly(A) tail length positively correlates with translational efficiency during the oocyte-to-embryo transition. **(A)** Correlation between poly(A) tail length and translational efficiency at each individual developmental stage determined by integration of our tail length data with published ribosome profiling data^47^. **(B)** Correlation between change in poly(A) tail length and change in translational efficiency between consecutive developmental stages, colored by change in tail length. **(C)** Poly(A) tail lengths of genes binned by translational efficiency (plot facets) at each individual developmental stage. Pairwise two-sided Wilcoxon tests shown for all stages compared to ZY (ns, p > 0.05; *, p <= 0.05; **, p <= 0.01; ***, p <= 0.001; ****, p <= 0.0001). Only stages with at least 10 genes in each tail length or translational efficiency bin are plotted. **(D)** Change in poly(A) tail length for polyadenylated genes with increase translational efficiency at each stage transition. Horizontal lines indicate mean. Pairwise two-sided Wilcoxon tests shown for all stage transitions compared to MII>ZY (****, p <= 0.0001).

A closer examination of the relationship between tail length and translation at each stage revealed stage-specific differences. Most strikingly, we found that for any given level of translational efficiency, mRNAs at the zygote stage were able to achieve the same level of translational efficiency with significantly shorter poly(A) tails compared to other stages (Fig. 5C. Consistent with this, significantly smaller increases in poly(A) tail length mRNAs were seen among mRNAs translationally activated from MII to ZY relative to all other stage transitions (Fig. 5D). Given that global mRNA tail lengths are shortest at the zygote stage, these findings raised the possibility that the tail length of an mRNA relative to other mRNAs a given stage is a greater determinant of translational efficiency than absolute poly(A) tail length.

### Global deadenylation reshapes relative mRNA poly(A) tail length distributions to activate translation prior to EGA

To further investigate the role of relative versus absolute tail length, we more closely examined the relationship between change in tail length and change in translational efficiency at each stage. Although different global trends in tail length are seen at different stage transitions (Fig. 1E-F), most mRNAs behave as predicted, with coordinated increases or decreases in both tail length and translational efficiency (Fig. 5B). However, during fertilization from MII to ZY, the global trend line was visibly shifted towards the upper left quadrant (Fig. 5B, MII to ZY), pointing again to a unique relationship between tail length and translational efficiency at the zygote stage. This shift was consistent with our findings above that show efficient translation of mRNAs with short tails--and with small increases in tail length--in the zygote relative to other stages (Figs. 5C-D). The shift further suggested the presence of a subset of mRNAs with an inverse relationship between translational efficiency and tail length—mRNAs with increased translational efficiency despite a decrease in tail length.

To quantify the scope and stage-specificity of this population of mRNAs with an inverse tail length-translational efficiency relationship, we independently examined mRNAs translationally activated and mRNAs deadenylated across each developmental transition. Indeed, mRNAs that were translationally activated but deadenylated were identified during both oocyte maturation and fertilization—the transitions marked by global deadenylation—but most pronounced from MII to ZY (Fig. 6A). In fact, between MII and ZY, 60% of mRNAs with increased translational efficiency demonstrated no accompanying increase in poly(A) tail length. Specifically, 43% of mRNAs were translationally activated despite significant decreases in tail length and an additional 17% demonstrated an increase in translational efficiency despite no significant change in tail length. In contrast, few translationally activated genes were deadenylated after the zygote stage (Fig. 6A). Similarly, examination instead of all mRNAs deadenylated during each transition showed that subsets of deadenylated mRNAs showed an increase in translational efficiency during both GV to MII and from MII to ZY (Fig. 6B). This trend, again, was most prominent across the transition from MII to ZY, during which almost 50% of deadenylated mRNAs also showed increased translation efficiency. From GV to MII, although the number of genes was relatively low, deadenylated mRNAs with increased translational efficiency were enriched for factors with important roles that reflect the coming switch to mitosis after fertilization, including “mitotic cell cycle phase transition,” “positive regulation of mitotic cell cycle,” “mitotic spindle organization,” and “mitotic sister chromatid segregation”). From MII to ZY, mRNAs deadenylated but translationally activated were enriched for roles in the early embryo, including “DNA-templated transcription, initiation”, “RNA splicing”, “mRNA processing”, and “regulation of DNA repair”. In contrast, deadenylated genes with decreased translational efficiency were enriched for genes involved in processes known to be downregulated during oocyte maturation (e.g. “cellular respiration,” “electron transport chain,” “ATP metabolic process,” “cytoplasmic translation”, and “ribosome biogenesis,”) or no longer needed after fertilization (e.g. “meiotic cell cycle”, “meiotic nuclear division,” “egg activation,” and “regulation of fertilization”) (Fig. 6C). This specificity, with respect both to stage and biological process, suggests translational activation of this subset of deadenylated mRNAs is a selective and regulated process.

**Figure 6.**
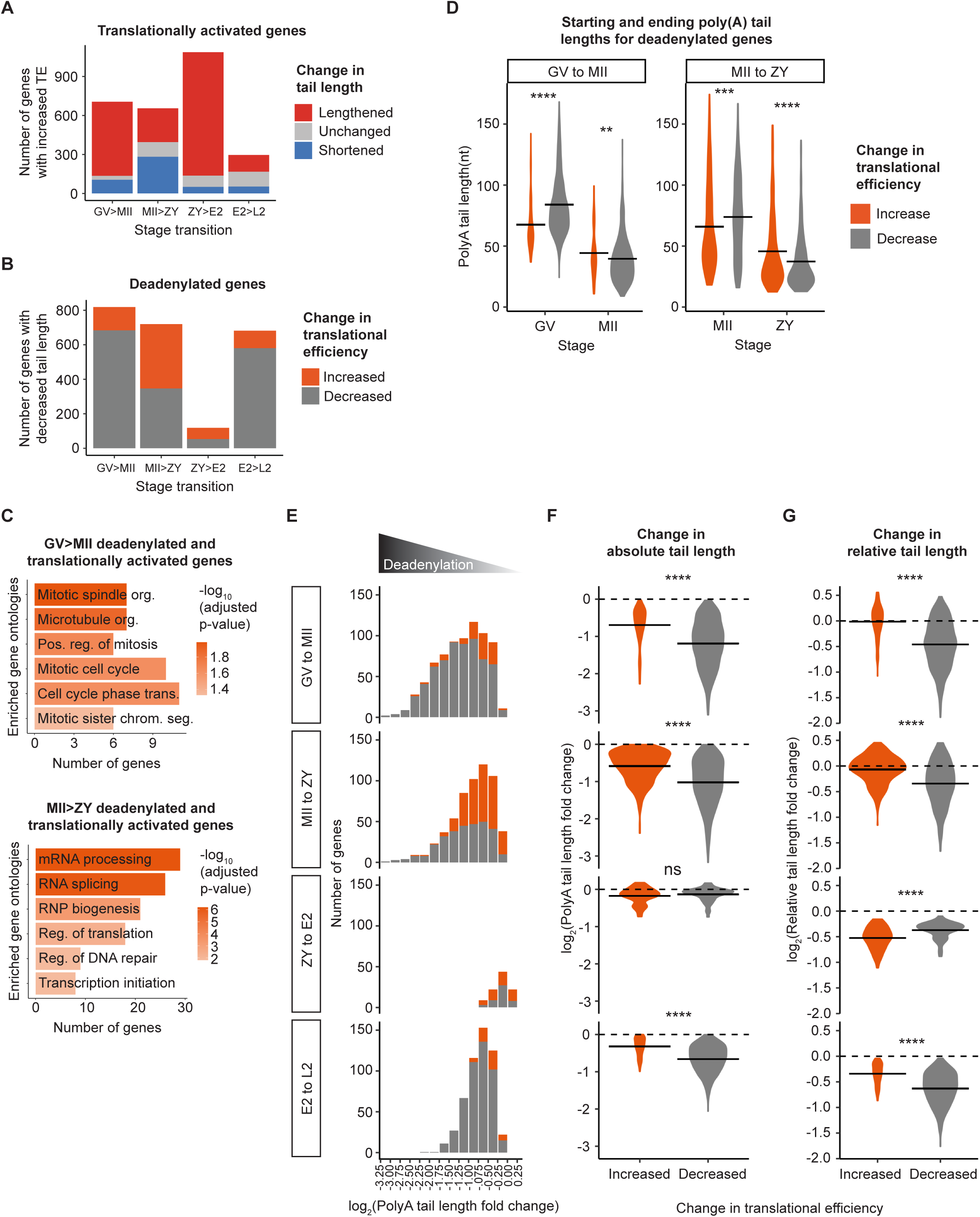
Global deadenylation reshapes relative mRNA poly(A) tail length distributions to activate translation prior to EGA. **(A)** Number of translationally activated genes with lengthened (red), shortened (blue), and not significantly changed tail length (grey). **(B)** Number of deadenylated genes with increased (orange) or decreased (grey) translational efficiency. **(C)** Enriched gene ontologies in genes deadenylated and translationally activated during oocyte maturation (GV>MII, top) or fertilization (MII>ZY, bottom). **(D)** Stage 1 and 2 geometric mean poly(A) tail lengths for deadenylated genes that are translationally activated (orange) or repressed (grey) during maturation (GV to MII) and fertilization (MII to ZY) stage transitions. Horizontal line indicates arithmetic mean. **(E)** Number of deadenylated genes that are translationally activated (orange) or repressed (grey), binned by magnitude of deadenylation. **(F-G)** Log2 fold change in absolute **(F)** or relative **(G)** tail length for deadenylated genes that are translationally activated (orange) or repressed (grey). Horizontal line indicates arithmetic mean. For **(B-G)**, to include genes translationally activated or repressed despite no significant change in tail length, the adjusted p-value cutoff for classifying genes as “deadenylated” was removed. Two-sided Wilcoxon tests shown (ns, p > 0.05; ****, p <= 0.0001).

To explore the mechanism by which this inverse relationship between change in tail length and translation might arise, we next compared tail length dynamics for deadenylated mRNAs that were translationally activated versus repressed. Interestingly, relative to mRNAs with decreases in translational efficiency, mRNAs with increased translational efficiency across both GV to MII and MII to ZY began each transition with significantly shorter tails, on average, but ended each transition with longer tails (Fig. 6D). Of note, although poly(A) tails for the activated genes were longer than that for the repressed genes, they were still relatively short overall with mean tail lengths of only 44 nucleotides and 46 nucleotides for those activated at GV to MII and MII to ZY, respectively.

These findings suggested that this population of translationally activated mRNAs—despite an overall mean decrease in tail length—retained longer tails relative to the translationally repressed mRNAs through relative protection from the global deadenylation occurring at these stages. Consistent with this idea, deadenylated mRNAs with increased translational efficiency during GV to MII and MII to ZY were highly enriched among those with the smallest decreases in tail length (Fig. 6E) and showed smaller decreases in absolute tail length compared to the repressed subset of deadenylated mRNAs (Fig. 6F). Further, we calculated each gene’s mean tail length relative to the global mean tail length at each stage, which we call “relative tail length” (see Methods), as a quantitative measure of the poly(A) tail length rank order. This comparison showed that deadenylated genes that were translationally activated exhibited a significantly smaller decrease in relative tail length than those that were translationally repressed (Fig. 6G). In fact, during both oocyte maturation and fertilization, the average change in relative tail length for the deadenylated genes that are translationally activated is close to zero, with a substantial subset of deadenylated genes that demonstrate an overall increase in relative tail length. In contrast, deadenylation does not appear to be a significant mechanism for translational activation after the zygote stage, pointing to a unique and transient shift in the translational regulatory regime in the late oocyte and early embryo. Together, these data support a model in which mRNA tail length rank-order can be reshaped by global deadenylation in a stage-specific manner to activate translation for specific subsets of mRNA—not by increases in absolute tail length—but by relative protection from deadenylation.

### Translational efficiency of maternal genes critical for development is regulated by dynamic changes in poly(A) tail length

In our analyses of all genes, we identified multiple mechanisms by which changes in tail length regulate mRNA translation and stability. To examine the potential importance of these specific mechanisms in development across the OET, we next asked whether maternal mRNAs with essential roles in development from oocyte to embryo demonstrate tail length dynamics consistent with regulation by these mechanisms. To do this, we examined poly(A) tail length dynamics for a group of 51 known maternal effect genes^61^ (MEGs) (Data S5). MEGs encode factors transcribed during oocyte growth for which the maternal (i.e., oocyte) contribution has been shown to play critical roles during the OET in oocyte and/or early embryo development^61–63^.

As observed amongst all genes, poly(A) tail lengths for this group of MEGs strongly correlate with translational efficiency at all five stages of the OET (Fig. 7A) and change in tail length and change in translational efficiency between consecutive developmental stages are also positively correlated (Fig. 7B). Examination of poly(A) tail dynamics of this group of mRNAs showed that the MEGs largely follow the global, stage-specific trends in poly(A) tail length that we observed for all genes (Fig. 1E-F). Specifically, as a group, MEG poly(A) tails are longest at GV with a mean tail length of 74 nucleotides, globally shortened during oocyte maturation, and further shortened during fertilization (Figs. 7C, S5A). Also similar to patterns observed for all genes, examination at the gene-specific level revealed specific subsets of MEGs at each stage transition for which regulation of mRNA tail lengths were opposite to these global trends as well (Figs. S5B-C). There were, however, two notable differences between tail length regulation for MEGs compared to the patterns of regulation we observed for all genes. First, at the gene-specific level, there was a trend towards MEGs being more likely to be polyadenylated during each developmental transition between GV to E2 relative to the rest of the transcriptome, particularly from ZY to E2 (Fig. 7D). In fact, virtually all MEG mRNAs demonstrated a significant increase in poly(A) tail length at one or more stages across the OET. This might be expected given the established importance of these genes for development during the OET. Second, poly(A) tails for MEGs were twice as likely to be deadenylated from E2 to L2 compared to all genes (Fig. 7D). Deadenylation of this subset of mRNAs is consistent with the expected clearance of a substantial proportion of maternal transcripts in association with major EGA.

**Figure 7.**
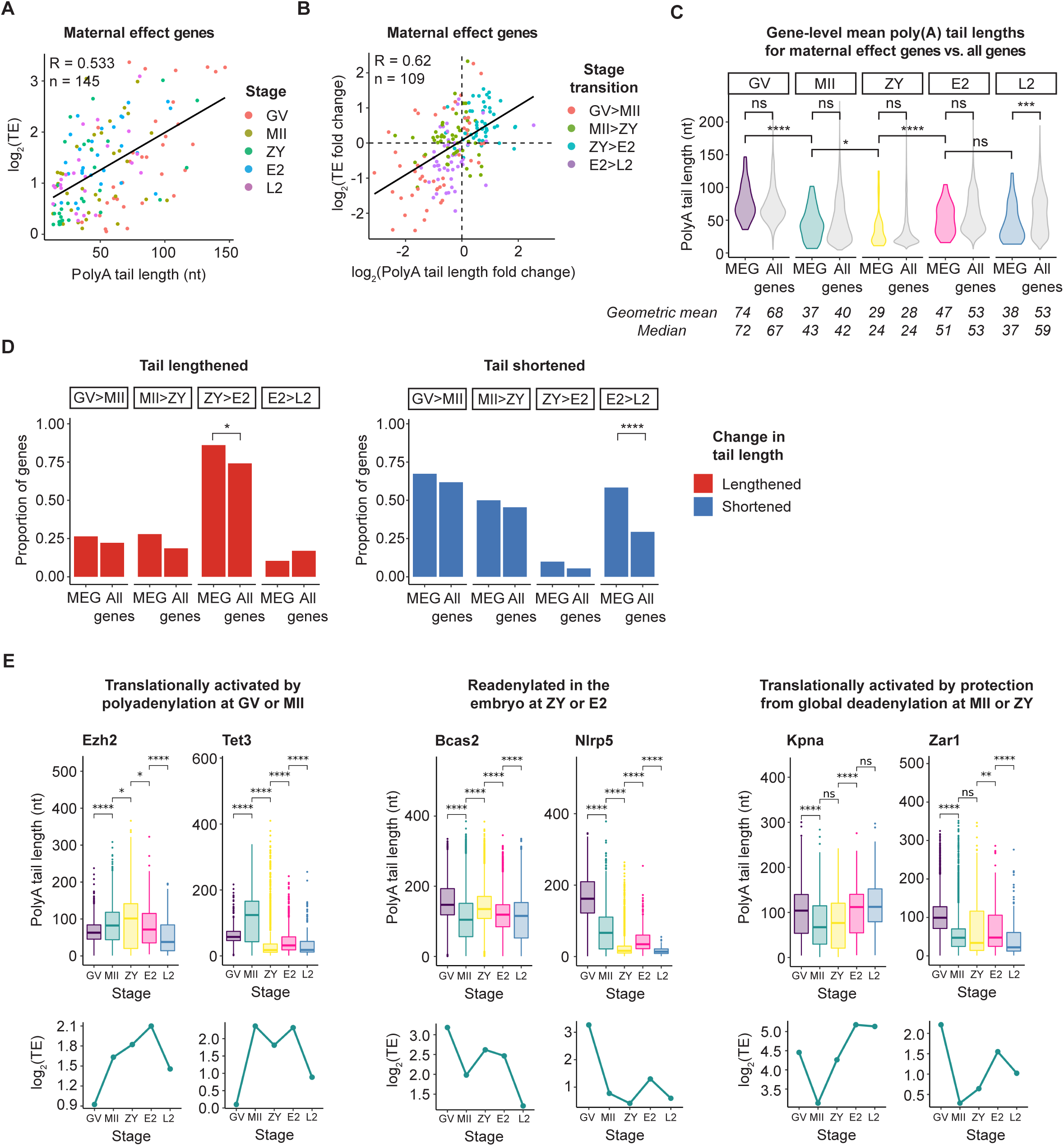
Translational efficiency of maternal effect genes (MEGs) critical for development is regulated by dynamic changes in poly(A) tail length. **(A)** Correlation between poly(A) tail length and translational efficiency for maternal effect genes. **(B)** Correlation between change in poly(A) tail length and change in translational efficiency between consecutive developmental stages for maternal effect genes. **(C)** Violin plots showing global distributions of gene-level mean poly(A) tail lengths at each stage transition for maternal effect genes (colored) compared to all genes (grey). Geometric means and medians of distributions given below violin plots. Only genes with a minimum of 10 polyadenylated reads in both replicates combined were included in this analysis. Pairwise two-sided Wilcoxon tests shown (ns, p > 0.05; *, p <= 0.05; ***, p <= 0.001; ****, p <= 0.0001). **(D)** Proportion of maternal effect genes or all genes with significantly (adjusted p-value < 0.05) lengthened (red, left) or shortened (blue, right) tail length at each consecutive developmental stage transition. Only maternal effect genes with at least 10 polyadenylated reads in both stages represented were included in this analysis. Pairwise one-sided Fisher’s exact tests shown (ns, p > 0.05; *, p <= 0.05; ***, p <= 0.001; ****, p <= 0.0001) (E) Poly(A) tail lengths (upper) and translational efficiencies (lower) of select maternal effect genes at each stage across the oocyte-to-embryo transition. Genes were filtered by a minimum of 1 FPKM in the RNA-seq dataset used to calculate translational efficiency. MEG = maternal effect gene; R = Pearson correlation coefficient; n = number of genes.

First, we asked whether a subset of MEGs demonstrated changes in tail length and translation efficiency consistent with selective, stage-specific cytoplasmic polyadenylation. Indeed, amid the global wave of deadenylation that characterizes the transition from GV to MII, 27% of the MEGs are significantly polyadenylated (Figs. 7D, 5B-C). Another 28% are significantly polyadenylated during the extended wave of deadenylation from MII to ZY although, as observed for all genes, mean increases in tail length at this stage are relatively small (Figs. 7D, 5B-C). Many of these MEGs play important roles in the zygote and early embryo, consistent with cytoplasmic polyadenylation of these MEGs being a selective process. Among these, *Trim28* is a scaffolding protein that recruits DNA methyltransferases and is critical for proper gene imprinting in the zygote and normal embryo development^64^. Consistent with its role in the zygote, *Trim28* mRNA is significantly polyadenylated during oocyte maturation, with a concurrent increase in translational efficiency (Fig. 7E). Similarly, the polycomb group protein *Ezh2*, which is critical for proper H3K9 and H3K27 methylation in the zygote^65^, is significantly polyadenylated from GV to MII and from MII to ZY, with associated increases in translational efficiency. In addition, the DNA demethylase *Tet3* is required for demethylation of the paternal genome following fertilization^66^, and consistent with this role, its mRNA demonstrates a transient increase in both poly(A) tail length and translational efficiency from GV to MII (Fig. 7E). A few additional examples are provided (Fig. S6A).

A second regulatory mechanism we observed in our analyses of all genes was uncoupling of deadenylation and decay during oocyte maturation (Fig. 3A), with readenylation of subsets of stable mRNAs in the next stage from MII to ZY and, even more prominently, during ZY to E2 (Figs. 4A-B, 4E-F). Examination of MEG tail dynamics provide further support for important biological roles for these mechanisms. With respect to deadenylation without decay during oocyte maturation, a greater proportion of MEGs deadenylated during oocyte maturation remain stable compared to all genes (61% of MEGs vs. 43% of all genes), and the large majority (70%) of these mRNAs also remain stable during fertilization. With respect to readenylation, MEGs are also enriched for genes identified as readenylated from MII to ZY (36% of deadenylated-stable for MEGs vs. 23% of deadenylated-stable for all genes), although, again, increases in tail length at this stage are small. As the maternal contribution of these MEGs is known to be critical for development, this tail lengthening likely represents readenylation of maternal transcripts as opposed to new transcription in the zygote. In support of this idea, none of these candidates for MEGs re-adenylated from MII to ZY overlap with predicted minor EGA genes^4^. As examples, *Bcas2* plays an important role in the DNA damage repair response in zygotes, and its depletion leads to arrest at the 2-cell to 4-cell embryo stage^67^. Deadenylated but stable during oocyte maturation, the poly(A) tail for *Bcas2* is significantly lengthened again between MII to ZY, with a concurrent increase in translational efficiency (Fig. 7E). Similarly, knockdown of homeobox protein *Sebox* results in arrest at the 2-cell stage through mechanisms that remain to be fully elucidated^68,69^. Consistent with its role in proper early embryo development, its mRNA remains stable despite being deadenylated during oocyte maturation and is subsequently significantly re-adenylated with concurrent increases in translational efficiency from MII to ZY (Fig. 7E). A few additional examples are provided (Fig. S6B).

Intriguingly, similar to the pattern seen for all maternal mRNAs, the vast majority of MEGs deadenylated but stable from GV to MII demonstrate significant tail lengthening during the first mitotic division in the embryo, with 90% (19 of 21) significantly polyadenylated between ZY and E2. This pattern in developmentally important genes underscores the potential role of readenylation at this stage (Fig. 4E-F). Although new transcription at this stage in the embryo might contribute to this observed increase in tail length, the maternally contributed products for these genes are known to be critical for development and only 2 genes (*Ctcf, Gclm*) overlap with >4000 genes previously predicted to be minor EGA genes^4^ and none overlap with a previously published list of >2500 genes predicted to be major EGA genes^55^. These data suggest that the increase in tail length from ZY to E2 for these MEGs predominantly represents readenylation of maternal transcripts. Examples of MEGs that are deadenylated during oocyte maturation and readenylated at E2 include *Nlrp5* (aka *Mater*), a critical component of the subcortical maternal complex (SCMC) whose absence results in 2-cell arrest^70,71^; *Nlrp9a*, another candidate component of the SCMC^72^; and *Pms2*, which plays an important role in DNA mismatch repair and in preventing replication errors in the early embryo^73^ (Fig. 7E). A few additional examples are provided (Fig. S6B).

Third, our analyses of all genes also uncovered subsets of mRNAs translationally activated despite decreases in mean tail length during the global waves of deadenylation from GV to MII and, even more notably, from MII to ZY (Figs. 6A-B). In the absence of detectable cytoplasmic polyadenylation, the proposed mechanism for translational activation of these genes is through increases in relative tail length as the tail rank order is reshaped transcriptome-wide during global deadenylation. To further examine this potential mechanism, we asked whether a subset of MEGs also demonstrates this inverse relationship between change in tail length and translation efficiency. Indeed, similar to our analyses of all genes, we find subsets of MEGs with tails that are shortened during oocyte maturation or fertilization but demonstrate a paradoxical increase in translation efficiency across the same stage transition. Also similar to our observations for all genes, this phenomenon was most pronounced from MII to ZY (21% of deadenylated genes during GV to MII vs. 44% during MII to ZY). We next asked if tail length for these MEGs is relatively protected during global deadenylation, as seen for all genes (Fig. 6D-F). Indeed, we observe the same trends. During both the GV to MII and MII to ZY transitions, deadenylated MEGs with increased translational efficiency demonstrate smaller decreases in both absolute and relative tail length compared to translationally repressed MEGs although statistical power was limited given the small number of genes (Fig. S5D-E). As an example, *Kpna6* (aka *Importin α7*) is a protein with roles in nuclear import and spindle assembly critical for early embryo development, with knockout leading to arrest at the 2-cell stage by a mechanism that remains unknown^74^. *Kpna6* mRNA is protected from deadenylation from MII to ZY and polyadenylated between ZY and E2 and demonstrates concurrent increases in translational efficiency across both transitions (Fig. 7E). Additional examples include *Rlim*, a ubiquitin ligase required for imprinted X chromosome inactivation in mouse embryos, and Zar1, an RNA binding protein required for proper division from 1-cell to 2-cell embryo stage. Both *Rlim* and *Zar1* mRNAs are deadenylated but demonstrate paradoxical increases in translational efficiency between MII and ZY (Fig. 7E). Additional examples are provided (Figs. S6C, see also *Rnf1*, *Spin1* and *Trim28* in S6A).

## Discussion

It was first recognized more than 30 years ago that regulation of maternal mRNA poly(A) tail length is an important mechanism controlling gene expression in the absence of transcription across the oocyte-to-embryo transition^12^. Recent technological advances have made it possible to determine mRNA poly(A) tail lengths transcriptome-wide across the entire OET in the small number of cells available for study mammals, including humans^44,75^. However, the full repertoire of poly(A) tail lengths across the OET in mice and how changes in poly(A) tail length across the OET affect translation across the entire OET in any mammal has remained unknown. Here, we determine and compare mRNA poly(A) tail lengths transcriptome-wide in five distinct stages across the transition from oocyte to embryo in mice. In addition, through integration of these data with published ribosome profiling data at the same developmental timepoints^47^, we demonstrate that tail length and translational efficiency are coupled across the entire transition, with polyadenylation and deadenylation directly linked to translational activation and repression, respectively, from the fully grown GV oocyte until after the onset of major EGA in the 2-cell mouse embryo. Our findings uncover regulation of mRNA poly(A) tail length at multiple levels, involving dynamic changes regulated at both the global and gene-specific level in a stage-specific manner. That these changes are matched by changes in translational efficiency provides the long-anticipated opportunity to identify candidates for factors and pathways essential for these earliest stages of development in mammals and to dissect the post-transcriptional mechanisms that control their expression. Although most of the regulatory principles are evolutionarily conserved, it is known that there are important differences in the developmental timing and the specific regulatory factors involved among organisms^1,76^. Therefore, a better understanding of the specific mechanisms in mammals is required both to advance our understanding of the earliest stages of our own development and to make meaningful improvements in the diagnosis and treatment of infertility. These datasets are a rich resource for the field and enable future studies of the post-transcriptional mechanisms linking poly(A) tail length, translation, and RNA stability to drive development across the transition from oocyte to embryo.

A surprise finding in this study was the marked wave of global tail shortening from MII to ZY. It has been known for decades that global deadenylation—with and without decay—is a prominent feature of the transition from GV to MII. However, we found that this wave of deadenylation continues during fertilization. This extended wave of global deadenylation results in accumulation of maternal mRNAs with the shortest tails across the transition at the ZY stage, with global mean tail length of 28 nucleotides (Figs. 1E-F). What might be the biological role for the continued and widespread tail shortening between MII and ZY? The roles are likely multiple and may include roles previously proposed for global deadenylation from GV to MII, including maternal mRNA clearance and deadenylation without decay to potentially facilitate reactivation or decay at later stages across the OET^2,26^. Global shortening of maternal mRNAs from MII to ZY might also allow preferential translation of newly transcribed embryonic transcripts during minor EGA. Newly transcribed, these mRNAs would be expected to have relatively long poly(A) tails^52–54^ while being low in abundance; therefore, global shortening of maternal mRNAs at this stage would potentially enhance translation of this small group of transcripts.

Our findings suggest an additional role for global deadenylation—and for the extended wave of deadenylation from MII to ZY—in the translational activation of specific subsets of maternal mRNAs with short tails. Decades of data support the current paradigm that translational activation is driven by cytoplasmic polyadenylation and increases in absolute poly(A) tail length. Here, however, we identify subsets of relatively short-tailed, maternal mRNAs translationally activated without detectable increases in poly(A) tail length (Fig. 6A). This phenomenon was observed during the transitions from GV to MII and MII to ZY—stages in which the global changes in tail length is dominated by deadenylation. We propose a model in which specific subsets of maternal mRNAs can be translationally activated during these waves of global deadenylation— not through increases in absolute tail length—but through relative protection from global deadenylation. In this model, the global decrease in tail length reshapes the rank order of mRNA tail lengths and shifts the minimum tail length required to achieve a certain level of translational efficiency, such that short-tailed mRNAs can be efficiently translated in the MII oocyte and zygote.

This model mechanism--and an important role for relative as opposed to absolute tail length--is supported by several lines of evidence. First, we find that mRNAs deadenylated but translationally activated undergo smaller decreases in both absolute and relative tail length than deadenylated mRNAs that are translationally repressed (Figs. 6E-G). As a result, this subset of mRNAs has tails that are overall shortened during global deadenylation but have a longer “relative tail length” after deadenylation compared to mRNAs that are translationally repressed at the same stage (Fig. 6G). Second, we find that mRNAs at the zygote stage were able to achieve the same level of translational efficiency with significantly shorter poly(A) tails compared to other stages (Fig. 5C). Third, the deadenylated mRNAs translationally activated at these stages are enriched for genes with roles specific for development of early embryo, further supporting the idea that this is a selective process and important for gene expression in the embryo (Fig. 6C). Finally, we find that even translationally activated mRNAs with significant increases in tail length demonstrate much smaller increases in tail length during MII to ZY relative to all other stage transitions (Fig. 5D). Collectively, these data suggest that relative poly(A) tail length is equally important as absolute tail length in translational activation at these early stages and, together, support a model in which global deadenylation reshapes the rank order of short-tailed maternal poly(A) tails to activate maternal mRNAs–not by increases in absolute mean tail length—but by increases in relative tail length by relative preservation of their tails in the face of global deadenylation.

Intriguingly, a similar mechanism has been described in Drosophila during egg activation, with translational activation of a subset of mRNAs with long tails in the setting of selective mRNA deadenylation without a requirement for cytoplasmic polyadenylation^43^. Although the details differ (e.g., long tails preserved during selective deadenylation in Drosophila versus short tails preserved during global deadenylation in mice), our findings suggest this mechanism is evolutionarily conserved and that deadenylation—and specifically, resistance to deadenylation— plays a critical, stage-specific role in activating translation during the OET. One potential role for this mechanism might be that, by significantly reducing the population of mRNAs with long tails during global deadenylation, mRNAs with short tails might better compete for a factor important for translation but of limited abundance at these early stages, such as PABP or ribosomes. In fact, it has been shown that PABP is present at limiting levels in Xenopus oocytes and that PABP abundance contributes to coupling and uncoupling of tail length and translation efficiency^58^. Intriguingly, oocytes express an oocyte-specific PABP called embryonic poly(A) binding protein (ePAB, aka PABPC1L)^77–79^. Whether the unique properties of ePAB might critically contribute to the relationship between poly(A) tail length and translational efficiency for short-tailed maternal mRNA at the MII and zygote stages is an interesting future question.

Another post-transcriptional mechanism of gene-regulation unique to the OET is the widespread uncoupling of deadenylation and mRNA decay. As previously mentioned, deadenylation is generally considered the first step in mRNA decay in somatic cells^19,20^. In fact, decay generally follows deadenylation so rapidly that decay intermediates are rarely captured. However, *in vitro* studies almost 50 years ago suggested that deadenylation and decay are uncoupled for a subset of mRNAs in the mouse during oocyte maturation^24–26^. Yet, it has remained unknown since this time what specific genes escape decay. Here, we observe two groups of deadenylated mRNAs at oocyte maturation—with large numbers of maternal mRNAs with decreases in abundance consistent with maternal mRNA decay as well as thousands that remain stable, consistent with uncoupling of deadenylation and decay for a large subset of maternal mRNAs (Fig. 3A). Intriguingly, we also observe similar groups of deadenylated mRNAs at later stages of the OET (Fig. 3A), suggesting that deadenylation and decay might remain uncoupled for a subset of mRNAs at stages after oocyte maturation. However, analyses of RNA abundance at these stages are confounded by new transcription, with the potential to overestimate the stable population, particularly from ZY to E2 and E2 to L2. Nevertheless, the presence of a similar trend was also seen from MII to ZY, where very little new transcription is predicted (Fig. 3A), supports the possibility that uncoupling continues to be an important mechanism regulating gene expression after oocyte maturation. As more sensitive RNA metabolic labeling approaches are developed, it will be important to differentiate maternally inherited and new embryonic transcripts more rigorously to dissect the relationship between deadenylation and decay in the early embryo.

As an additional role for global deadenylation, we propose that deadenylation uncoupled from decay during this extended wave of global deadenylation from GV to ZY also acts to broadly prime maternal mRNAs for readenylation and translational activation in the embryo. In support of this model, we identify a subset of mRNAs deadenylated but stable from GV to MII with evidence of tail lengthening again from MII to ZY (Fig. 4A-B, MII>ZY). Multiple findings suggest this tail lengthening represents readenylation of maternal transcripts: (i) there is minimal new transcription at this stage anticipated from MII to ZY, (ii) there is minimal overlap of these potential readenylated genes with those predicted to be transcribed during minor EGA^4^, (iii) the readenylated mRNAs are highly associated with ribosomes and enriched for mRNAs with increased translation efficiency (Fig. 4C, MII>ZY) and (iv) the readenylated mRNAs are enriched for genes encoding factors with important roles specific for the new embryos (Fig. 4D). This pattern of tail regulation is even more striking during the first embryo cleavage. For >85% of mRNAs deadenylated without decay during oocyte maturation, readenylation occurs not during fertilization, but later during the transition from ZY to E2 (Fig. 4A-B, ZY>E2). In fact, the bulk of maternal mRNAs—including MEGs—are polyadenylated and translationally activated during this short, 6-hour transition from ZY to E2, marking a major switch in tail regulation during this developmental window (Figs. 2B, 7D). Together, our findings suggest global deadenylation in the oocyte primes maternal mRNAs for readenylation in the embryo and plays a major role in driving the shift from decreased to increased tail length observed from ZY to E2 to regulate translation across the OET (Figs. 2B, 4A-E, ZY>E2).

Despite the exceptions observed and discussed here, the overall relationship between mRNA poly(A) tail length and translation across the transition from oocyte to embryo in mammals fits the paradigm that has been in place for decades, with parallel increases and decreases in tail length and translation efficiency. In fact, even during global deadenylation, we observe specific subsets of mRNAs with patterns consistent with cytoplasmic polyadenylation in the face of this wave (Figs. 2A-B). Yet, despite decades of study, this relationship remains poorly understood, particularly in mammals. These transcriptome-wide datasets provide an exciting opportunity to identify patterns in tail length and translation regulation to untangle the complex, stage-specific, post-transcriptional regulatory mechanisms controlling gene expression across the OET. Specific 3’UTR sequence motifs and the RNA binding proteins that bind them are known to play major roles^12,29,80–82^. The most well-studied of these is the 3’UTR cytoplasmic polyadenylation element (CPE) and the cytoplasmic polyadenylation element binding protein 1 (CPEB1), which play critical roles in regulating both deadenylation and cytoplasmic polyadenylation of maternal mRNAs during oocyte maturation^12,15^. However, CPEB1 becomes undetectable after fertilization, and while important inroads have been made, the 3’UTR motifs required to regulate poly(A) tail length in the embryo before EGA are less explored^29^. Important next steps include motif analyses to identify the known and novel 3’UTR motifs—and combinations of motifs—that drive stage-specific developmental events in the early mammalian embryo.

In addition to these RNA regulatory mechanisms, these data also provide an important opportunity to identify novel candidates for factors with important developmental roles at each stage and to order expression of these factors over developmental time from oocyte to embryo. In support, we show that genes polyadenylated at each stage are enriched for factors known to play key roles in important and temporally relevant developmental processes (Fig. 2C). As further validation, we examine patterns of tail regulation for mRNAs encoding a group of previously identified MEGs—maternal effect genes already known to be important drivers of development during the OET^61^. Even within this small group of genes, we find multiple examples of mRNAs with patterns of regulation consistent with each of the mechanisms described here—including mRNAs with tail lengthening consistent with stage-specific cytoplasmic polyadenylation, mRNAs deadenylated without decay during global deadenylation in the oocyte and readenylated at later stages in the embryo, and mRNAs with short tails translationally activated without detectable polyadenylation during waves of global deadenylation. Many of these factors are required for major EGA^61^, and as ∼50% of human embryos arrest at major EGA^7^, are candidates for factors important for female fertility.

Altogether, our findings demonstrate the presence of an extended wave of global deadenylation that sets up a switch in translation control across the transition from oocyte to embryo. Maternal mRNA tail length and translation efficiency is dynamically shaped by deadenylation and resistance to deadenylation in the oocyte and by broad readenylation of deadenylated-stable maternal mRNAs in the embryo. These divergent mechanisms in the oocyte and embryo support a model in which the tail length of an mRNA relative to other mRNAs a given stage is a greater determinant of translational efficiency than absolute poly(A) tail length. The datasets provided here offer a rich resource and exciting insight into the complex post-transcriptional mechanisms by which cytoplasmic polyadenylation and deadenylation shape and reshape the landscape of poly(A) tails to drive gene expression during the earliest stages of mammalian development.

## Materials and Methods

### Oocyte and embryo isolation

Mouse oocytes were isolated from C57B6J female mice. For GV oocytes, 4-week old females were stimulated by intraperitoneal injection of 5 IU pregnant mare serum gonadotropin (PMSG; Lee BioSolutions, 493-10), ovaries were harvested 48 hours after PMSG injection, and denuded oocytes collected after brief digestion with trypsin (Fisher Scientific, 25200056), collagenase (Sigma-Aldrich, C9407), and DNAse in M2 media (Sigma-Aldrich, M7167). For MII oocytes, mice were injected with 5 IU HCG (Sigma-Aldrich, C1063) 48 hours after PMSG induction and ovulated MII oocytes collected from the ampulla 15 hours post-HCG and cumulus cells by brief hyaluronidase (Sigma-Aldrich, H4272) digestion per standard protocol. Mouse embryos were isolated from B6D2F1/J female mice mated with C57B6J males of known fertility at the time of hCG injection. 1-cell, early 2-cell, and late 2-cell embryos were collected at 27-28, 30-32, and 46-48 hrs post-HCG.

### RNA extraction

For each stage and replicate, total RNA was isolated from ∼400 oocytes or embryos with Trizol reagent (Fisher Scientific, 15-596-018), followed by ethanol precipitation and RNA Clean & Concentrator-5 kit (Zymo Research, R1014) as per manufacturer’s protocol. RNA was eluted in 7 ul nuclease-free water and used immediately for Nanopore PCR-cDNA sequencing library prep.

### Design of polyadenylated standards

cDNA standards with the following sequences were obtained from IDT: ***pA_cDNA_standard_10:*** CTTGCCTGTCGCTCTATCTTCTTTTTTTTTTACCAACGGCGACGAATAGTAGTTTACTTCCTCCCTGCGGGCCCCTCCTGAAGTGCCACCTATACGGCTTGTTGAAGCCGATTAGTACAATAGATTTATTCAACCCCAAAGGTCTACACTCCCGGCTTACTCTTAGCTGATATGTCGCGCAATATCAGCACCAACAGAAA

#### pA_cDNA_standard_40

CTTGCCTGTCGCTCTATCTTCTTTTTTTTTTTTTTTTTTTTTTTTTTTTTTTTTTTTTTTTCGGCA AAGAGACAATTATAGCGGCTAGGAACGCAACTAGTTATAACGAACGGCCTCGAATAGTAGAAAATATCCCTCCTCCGGGCACCTCCTGAAATGCCACATATTCGGGTTATTGGCAATATCAG CACCAACAGAAA

#### pA_cDNA_standard_50

CTTGCCTGTCGCTCTATCTTCTTTTTTTTTTTTTTTTTTTTTTTTTTTTTTTTTTTTTTTTTTTTTTTTTTAGAGAGCCAGCAACAATTGCAAATGTCAGATCAAAGTAATATTAGCAAACAATAAGTCCCTAACTAGTTGTGACCTTTTGTAAAGTGAATTTCATTATATATGCTGTGCAATATCAGCACCAACAGAAA

#### pA_cDNA_standard_70

CTTGCCTGTCGCTCTATCTTCTTTTTTTTTTTTTTTTTTTTTTTTTTTTTTTTTTTTTTTTTTTTTTTTTTTTTTTTTTTTTTTTTTTTTTGCAACGGGGAGCCGAGATTATTGAGTGATCACCAGTAGCTGTACTATTATATAAGCTATTAAAGATTGATCAAAGTAAACATACCGCGCAATATCAGCACCAACAGAAA

#### pA_cDNA_standard_100

CTTGCCTGTCGCTCTATCTTCTTTTTTTTTTTTTTTTTTTTTTTTTTTTTTTTTTTTTTTTTTTTTTTTTTTTTTTTTTTTTTTTTTTTTTTTTTTTTTTTTTTTTTTTTTTTTTTTTTTTCCCAAGCGAAAACGGGTGCGTGGACTAGCGAGGAGCAAACGAAAATTCTTGGCCTGCGCAATATCAGCACCAACAGAAA

#### pA_cDNA_standard_130

CTTGCCTGTCGCTCTATCTTCTTTTTTTTTTTTTTTTTTTTTTTTTTTTTTTTTTTTTTTTTTTTTTTTTTTTTTTTTTTTTTTTTTTTTTTTTTTTTTTTTTTTTTTTTTTTTTTTTTTTTTTTTTTTTTTTTTTTTTTTTTTTTTTTTTAGGAGGCAATTCTATAAGAATGCATACGCAATATCAGCACCAACAGAA A

#### pA_cDNA_standard_150

CTTGCCTGTCGCTCTATCTTCTTTTTTTTTTTTTTTTTTTTTTTTTTTTTTTTTTTTTTTTTTTTTTTTTTTTTTTTTTTTTTTTTTTTTTTTTTTTTTTTTTTTTTTTTTTTTTTTTTTTTTTTTTTTTTTTTTTTTTTTTTTTTTTTTTTTTTTTTTTTTTTTTTTTTTCGAGCACGCAATATCAGCACCAACAGAAA

### Nanopore PCR-cDNA sequencing library prep

Approximately 400 oocytes or embryos per biological replicate per stage were used per sequencing run. An identical amount of ERCC spike-in standards^51^ per cell was added to samples at the beginning of RNA extraction. Nanopore PCR-cDNA sequencing (PCS) was performed with an early access cDNA-PCR Sequencing Kit (PCS-110) as per manufacturer’s protocol (Fig. S1A). This kit is most similar to the currently available cDNA-PCR Sequencing Kit (PCS-111). Briefly, ligation of Nanopore’s proprietary cDNA-RT adapter (CRTA) to the 3’ end of polyadenylated mRNAs enriched for transcripts with a minimum polyA tail length of 10 nt. Following enzymatic digestion of the adapter, RT with strand switching was performed, followed by selection of full-length transcripts by PCR amplification for 18 cycles. Sequencing adapters are then attached, followed by loading of the library onto a R9.4.1 flow cell (FLO-MIN106D) and sequenced in a Minion sequencing device with Minknow software (version 22.05.5). Libraries comprised of polyadenylated cDNA standards were prepared by PCR amplification using the following barcoded PCR primers from the Nanopore PCR-cDNA Barcoding Kit (SQK-PCB109), followed by attachment of sequencing adapters from the PCS-110 kit and sequencing on the Minion as per the PCS-110 protocol: pA_cDNA_standard_10: BC01, pA_cDNA_standard_40: BC02, pA_cDNA_standard_50: BC03, pA_cDNA_standard_70: BC04, pA_cDNA_standard_100: BC05, pA_cDNA_standard_130: BC06, and pA_cDNA_standard_150: BC07.

### Data processing

Raw fast5 files were basecalled with Guppy (version 6.0.6) following parameters: --config dna_r9.4.1_450bps_hac.cfg --fast5_out --trim_strategy none --records_per_fastq 0 --device cuda:all:100% --recursive. and full length reads with all adapter sequences (from the CRTA, PCR primers, and sequencing adapter) were filtered by Pychopper (version 2.5.0) following parameters: -k PCS110. Only full-length reads with a minimum basecalling Q-value score of 9 and all adapter sequences were retained for further analysis. Reads were mapped to the GRCm38 mouse transcriptome with minimap (version 2.17) using the following parameters: -a -x map-ont. Reads per transcript were quantified with Salmon (version 0.14.1) using the following parameters: salmon quant -l A --noErrorModel.

### Tail length measurements and comparisons

PolyA tail lengths were measured using tailfindr (version 1.3) using default parameters. Since tail length measurements correlated highly between replicates of the same stage, polyadenylated reads from the same stage were combined for further analyses. Gene-level polyA tail lengths were estimated as the geometric mean of all polyadenylated reads detected. For consecutive stage comparisons at the gene-level, genes were filtered by a minimum number of 10 polyadenylated reads in each of the stages compared. Significant changes in tail length were defined by a maximum (Benjamini-Hochberg) adjusted p-value cutoff of 0.05.

### PCA and Hierarchical clustering

Unsupervised hierarchical clustering of stages and biological replicates by poly(A) tail length was performed in R (version 4.1.2) with the pheatmap package (version 1.0.12). Principal component analysis (PCA) by poly(A) tail length was performed using the prcomp function in the stats package (version 4.1.2) and PCA by gene expression was performed using the plotPCA function in DESeq2 (version 1.34.0).

### Differential expression analysis

Read counts were normalized with ERCC standards by dividing read counts for each gene by the mean read count amongst ERCCs. Differential expression analysis was performed with DESeq2 (version 1.34.0) and differentially expressed genes (DEGs) were defined by a minimum fold change of 2x and those that did not meet this criteria were classified as “stable.”

### Translational efficiency analyses

Translational efficiencies (TEs) were obtained from a previously published ribosome profiling and RNA-seq dataset^47^ and TE was calculated as (FPKM + 1/FPKM + 1) as per the authors. Unless otherwise indicated, for comparisons of gene-level polyA tail length and translational efficiency, genes were filtered by a minimum of 25 FPKM in the RNA-seq dataset and 10 polyadenylated reads in our PCS dataset.

### Relative tail length analyses

Tail lengths relative to the stage-specific global mean (aka “relative tail lengths”) were calculated for each gene as follows: *(TL_rel_)_gene X_ = (TL_abs_)_gene X_* - (*mean TL)_stage Y_ + 70 nt*, where (TL_rel_)_Gene X_ is the relative tail length for a given gene X; (TL_abs_)_Gene X_ is its absolute tail length; and (mean TL)_Stage Y_ is the mean of mean tail lengths amongst all genes at stage Y. Note that all means were geometric means and that the scalar of 70 nt was added to avoid taking the log of negative numbers.

### Gene ontology overrepresentation analysis

Gene ontology overrepresentation analysis was performed with the Bioconductor package clusterProfiler (version 4.2.2) using gene ontology biological process annotations. Significantly enriched gene ontologies were determined by a Benjamini-Hochberg adjusted p-value cutoff of < 0.05.

### Data availability

The Nanopore PCR-cDNA sequencing data associated with this study will be made publicly available on GEO.

---

**Figure S1.**
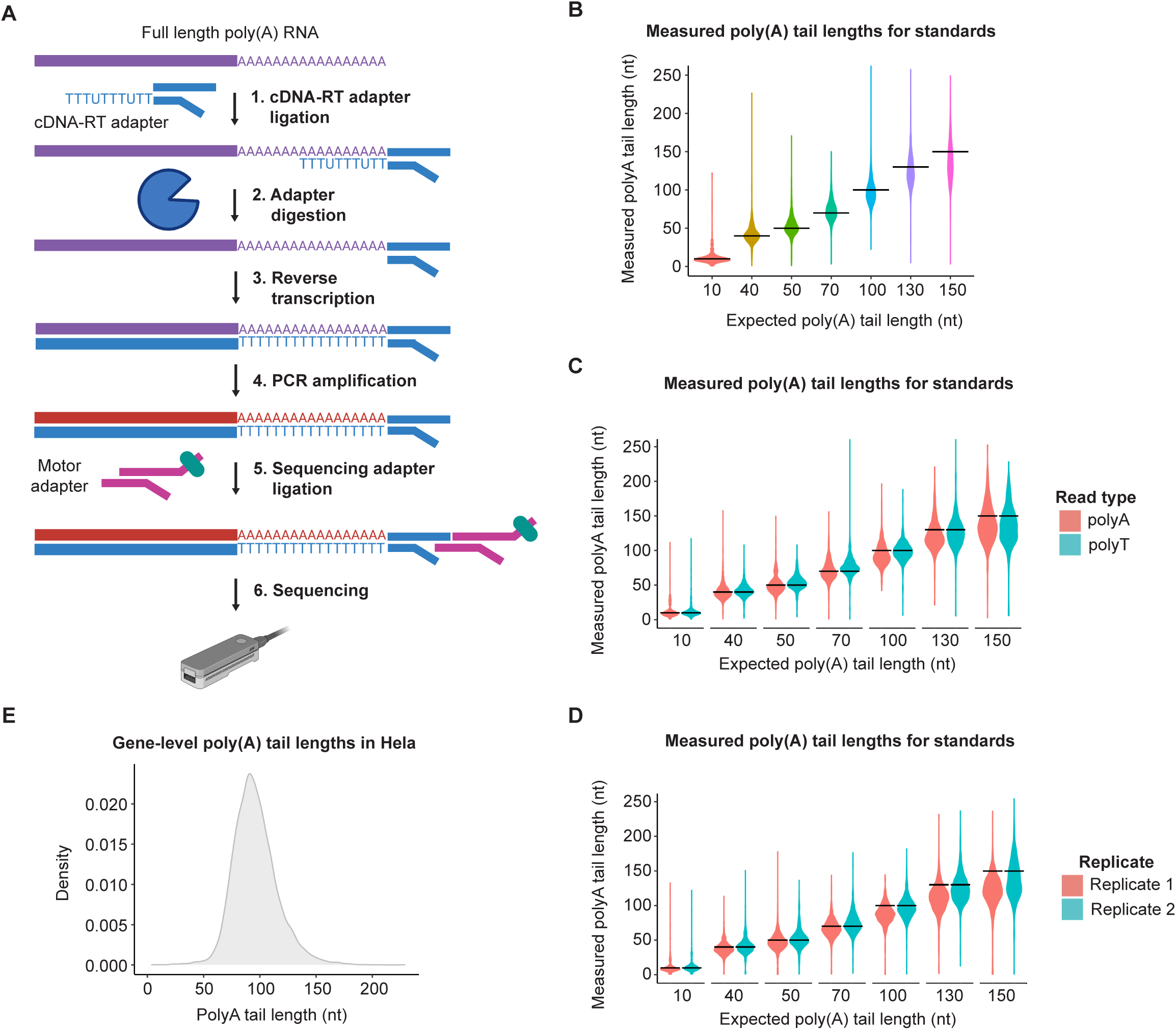
Nanopore PCR-cDNA sequencing accurately and reproducibly measures poly(A) tail lengths. **(A)** Schematic of Nanopore PCR-cDNA sequencing. **(B)** Measured tail lengths for 7 standards of different poly(A) tail lengths. **(C-D)** Same as **(B)** but separated by read type **(C)** or replicate **(D)**. For **(B-D)**, horizontal lines indicate expected poly(A) tail lengths for each standard. **(E)** Global distribution of gene-level mean poly(A) tail lengths in Hela. Only genes with a minimum of 10 polyadenylated reads were included in this analysis.

**Figure S2.**
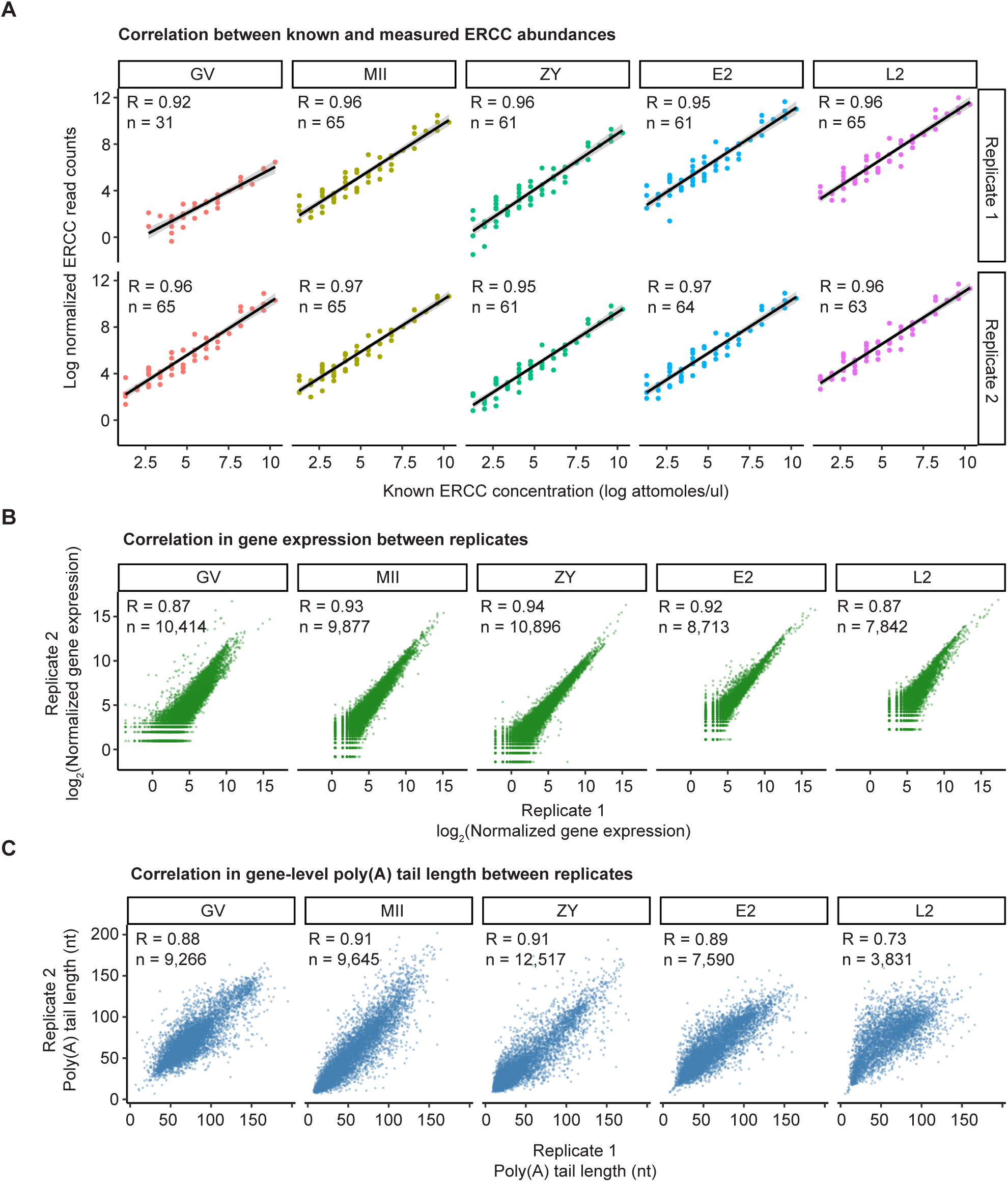
Nanopore PCR-cDNA sequencing reproducibly captures poly(A) tail lengths and gene expression profiles across the oocyte-to-embryo transition. **(A)** Correlation between known and measured abundances for ERCC standards. **(B)** Correlation in gene expression between replicates. **(C)** Correlation in gene-level poly(A) tail length between replicates. Only genes with a minimum of 20 reads with measurable poly(A) tail lengths in each replicate are plotted. R = Pearson correlation coefficient. N = Number of genes or ERCC standards.

**Figure S3.**
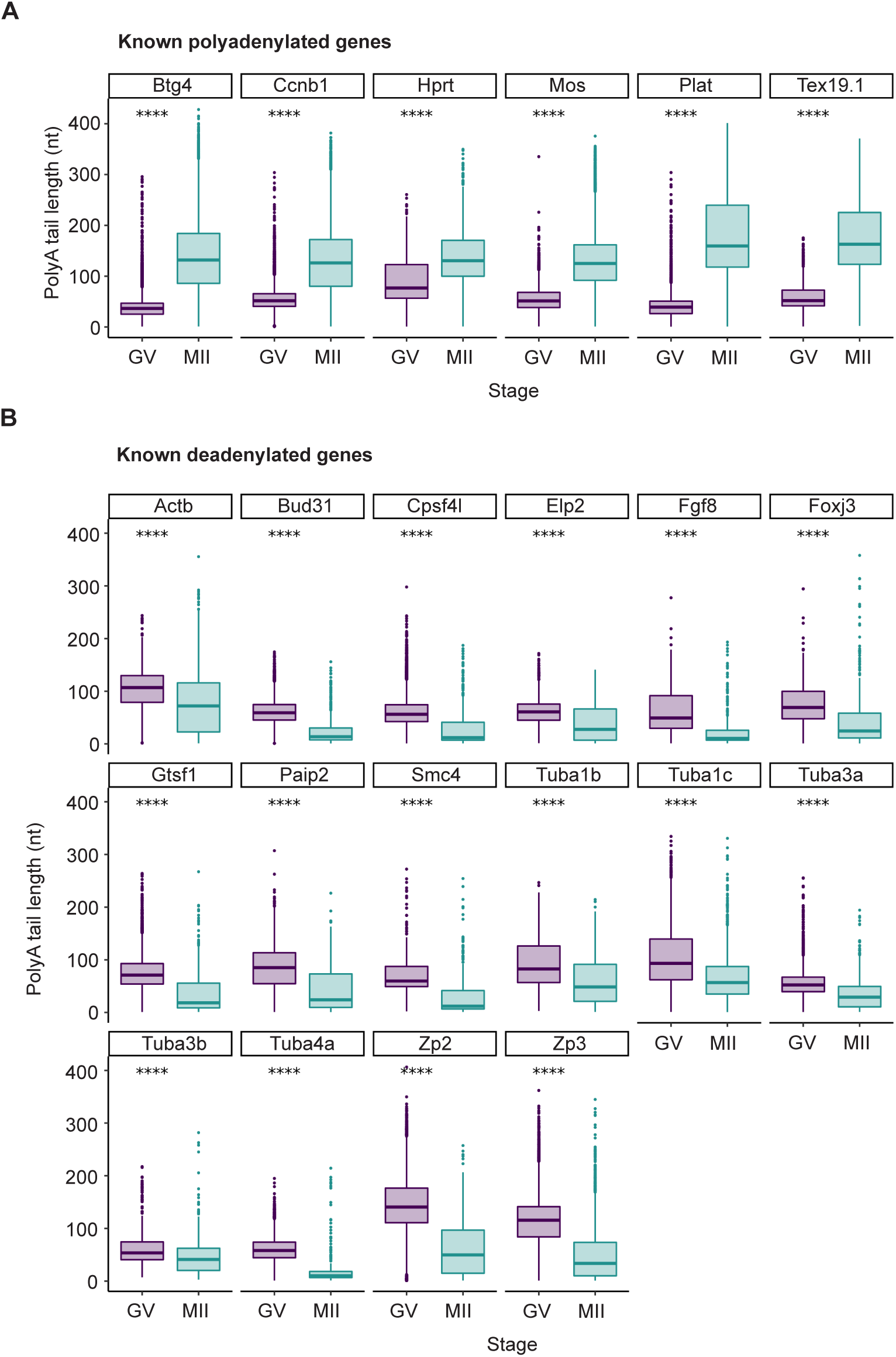
PCR-cDNA sequencing accurately captures known changes in poly(A) tail length for individual genes. **(A&B)** Measured poly(A) tail lengths in GV and MII oocytes for individual genes known to be polyadenylated **(A)** or deadenylated **(B)** during oocyte maturation. One-sided Wilcoxon tests shown (**** p <= 0.0001).

**Figure S4.**
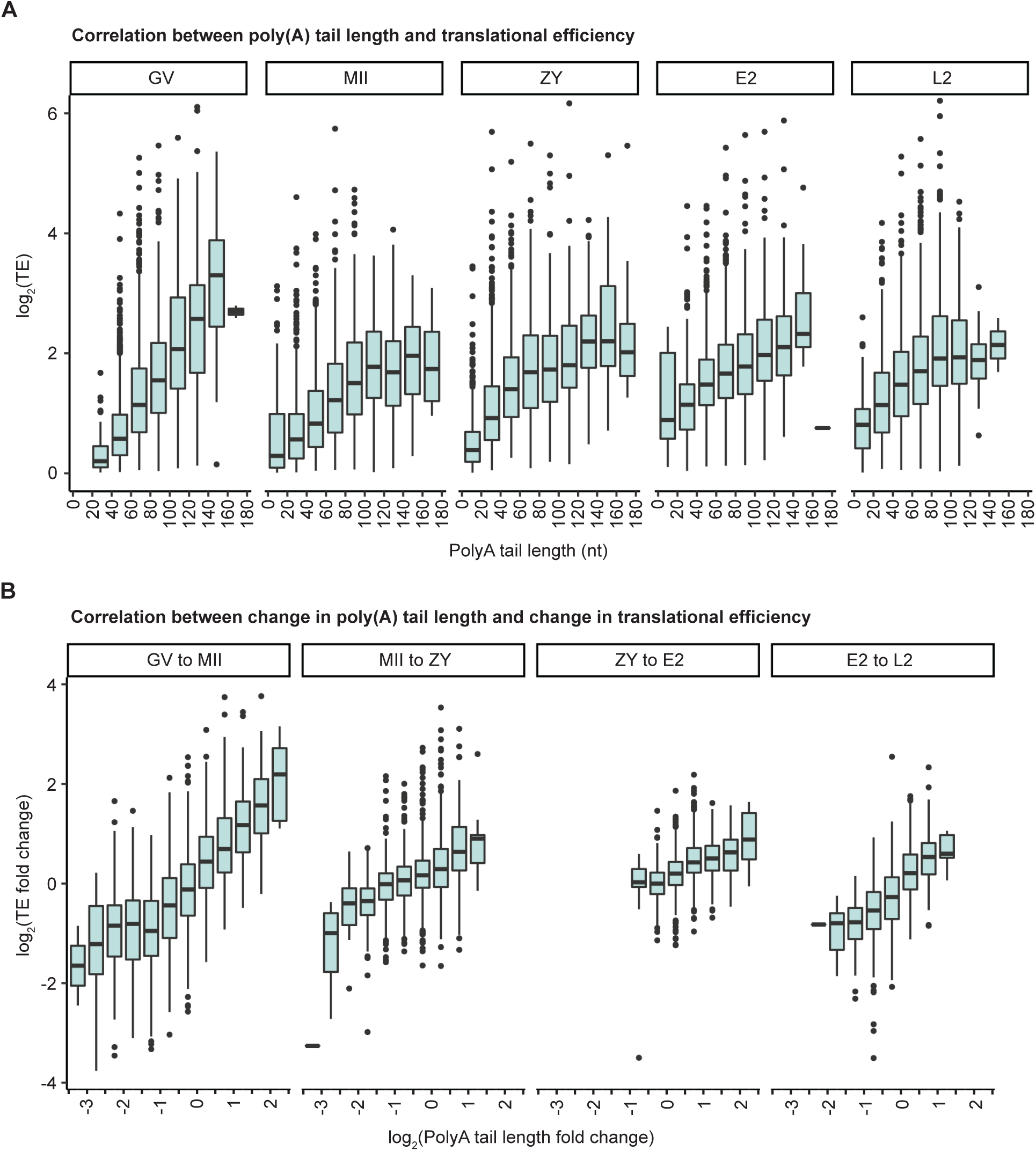
Poly(A) tail length positively correlates with translational efficiency during oocyte-to-embryo transition. **(A)** Translational efficiency of genes binned by poly(A) tail length at each developmental stage. **(B)** Change in translational efficiency of genes binned by change in poly(A) tail length between consecutive developmental stages.

**Figure S5.**
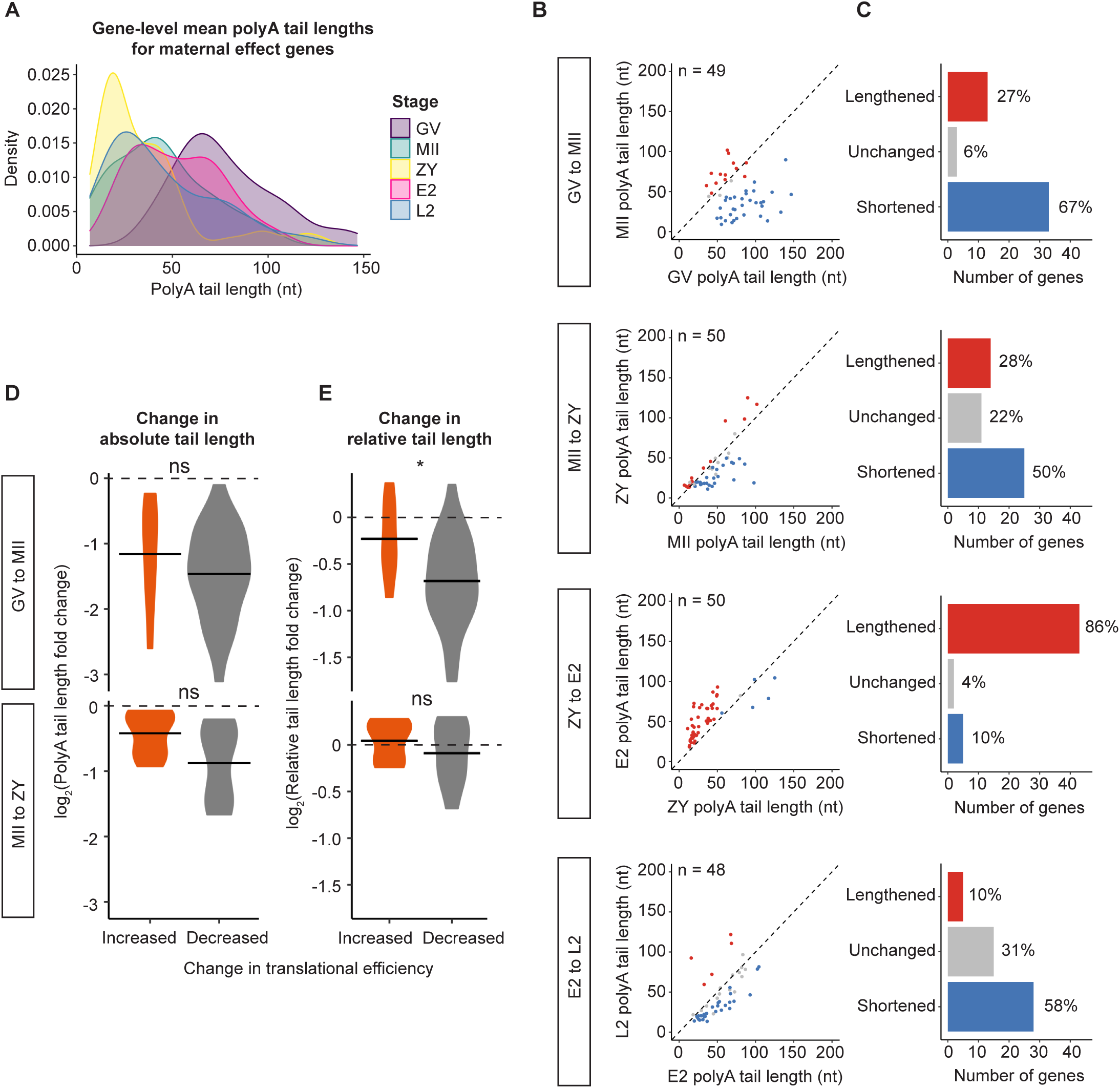
Dynamic regulation of poly(A) tail length of maternal effect genes. **(A)** Density plots showing global distributions of gene-level mean poly(A) tail lengths across the oocyte-to-embryo transition for maternal effect genes. **(B)** Scatter plots showing geometric mean poly(A) tail lengths for maternal effect genes with significantly (adjusted p-value < 0.05) increased (red), decreased (blue) or not significantly changed (grey) tail length at each consecutive developmental stage transition. Only genes with at least 10 polyadenylated reads in both stages represented were included in this analysis. **(C)** Number of genes shown in **(B)**. **(D-E)** Log2 fold change in absolute **(D)** or relative **(E)** tail length for deadenylated maternal effect genes that are translationally activated (orange) or repressed (grey). Horizontal line indicates arithmetic mean. Two-sided Wilcoxson tests shown (ns, p > 0.05; *, p <= 0.05).

**Figure S6.**
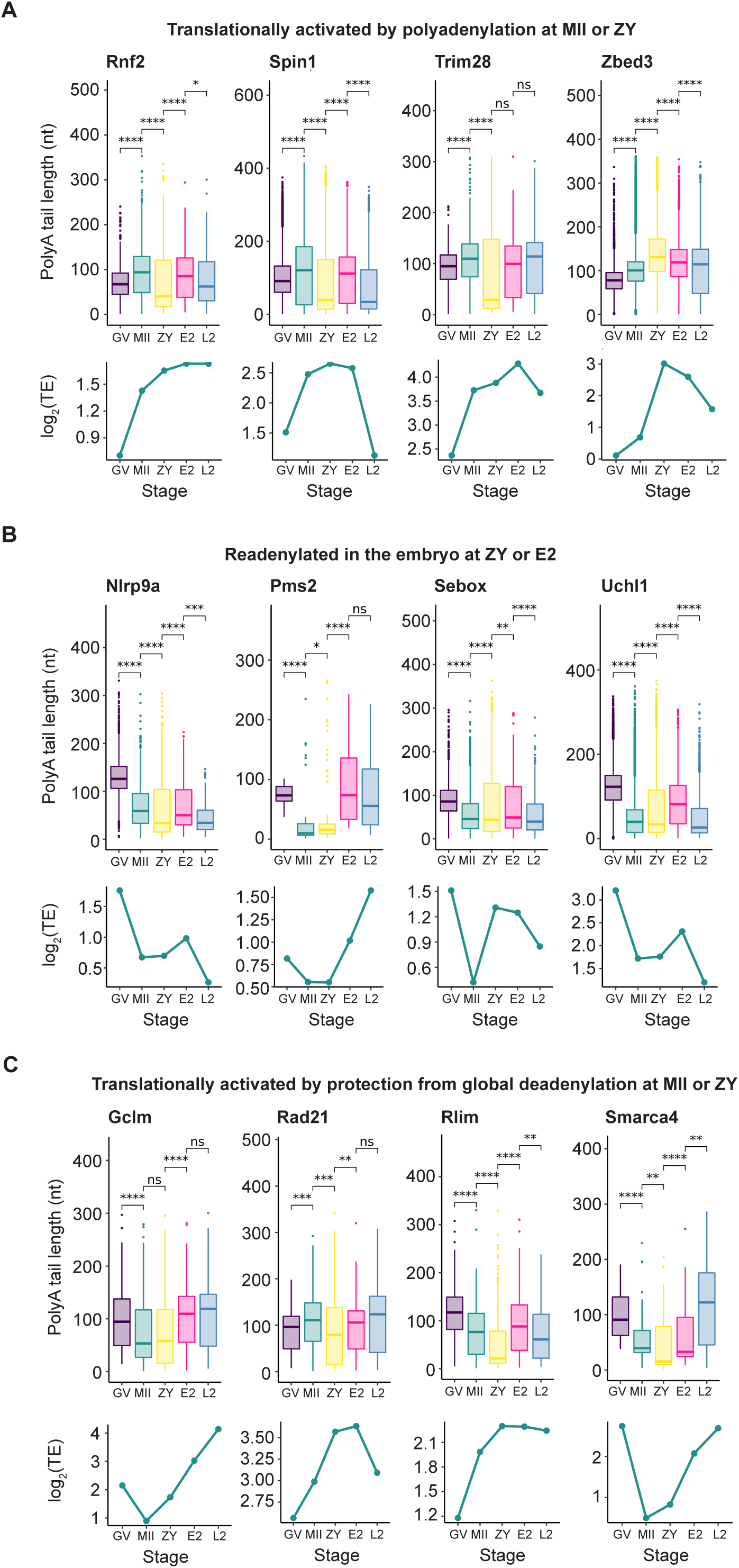
Example maternal effect genes. **(A-C)** Poly(A) tail lengths (upper) and translational efficiencies (lower) of select maternal effect genes across the oocyte-to-embryo transition.

